# Improved multimodal protein language model-driven universal biomolecules-binding protein design with EiRA

**DOI:** 10.1101/2025.09.02.673615

**Authors:** Wenwu Zeng, Haitao Zou, Xiaoyu Li, Yutao Dou, Xiaoqi Wang, Shaoliang Peng

## Abstract

The interactions between proteins and biomolecules form a complex system that supports life activities. Designing proteins capable of targeted biomolecular binding is therefore critical for protein engineering and gene therapy. Here, we propose a new generative model, EiRA, specifically designed for universal biomolecular-binding protein design, which undergoes two-stage post-training, i.e., domain-adaptive masking training and binding site-informed preference optimization, based on a general multimodal protein language model. A systemic evaluation reveals the SOTA performance of EiRA, including structural confidence, diversity, novelty, and designability on 8 test sets across 6 biomolecule types. Meanwhile, EiRA provides a better characterization for biomolecular-binding proteins than generic model, thereby improving the predictive performance of various downstream tasks. We also mitigate severe repetition generation in the original language model by optimizing training strategies and loss. Additionally, we introduced DNA information into EiRA to support DNA-conditioned binder design, further expanding the boundaries of the design paradigm. Purification experiments and molecular dynamics simulations verified the manufacturability and DNA-binding ability of the designed highly differentiated protein. Remarkably, EiRA achieved the “one-shot” design of a Glucagon peptide binder with SPR-confirmed micromolar affinity.

## Introduction

Over hundreds of millions of evolutionary years, proteins have emerged as the primary molecular mediators of biological processes. As of 28 March 2025, the UniParc database ^1^ archives 916,871,247 non-redundant protein sequences, yet this constitutes a vanishingly small fraction of the theoretically accessible sequence space. The engineering of novel proteins with tailored functionalities holds transformative potential for biomedical innovation, particularly in therapeutic development like immunotherapy and gene editing. Conventional approaches, including energy minimization-based rational design and physics-driven modeling, however, struggle to efficiently navigate this vast combinatorial landscape. Artificial Intelligence (AI)-enabled high-throughput design paradigms now promise to overcome this bottleneck. Post-2022, AI-generated protein candidates have surged at an unprecedented rate, outpacing the cumulative progress of the preceding decade, a testament to the catalytic role of AI in reshaping protein design paradigms ^2-4^.

Contemporary generative AI-driven *de novo* protein design frameworks typically follow a three-stage pipeline: (1) Functionally oriented backbone generation prioritizes structural stability and designability through geometric constraints derived from target motifs (e.g., binding interfaces or catalytic sites) or target biomolecular, employing probabilistic sampling via diffusion models (RFdiffusion ^5^, Chroma ^6^, and SCUBA-D ^7^) or flow-matching (Proteína ^8^) to explore conformationally diverse scaffolds or binders; (2) Spatial-aware sequence design tools such as ProteinMPNN ^9^, CarbonDesign ^10^, MapDiff ^11^, and ESM-IF1 ^12^ then populate amino acid residues onto these backbones using the geometry-aware neural networks to encode atomic-level spatial constraints, followed by autoregressive decoding to optimize sequence-structure compatibility; (3) Computational validation via protein structure predictors like AlphaFold2/3 ^13, 14^ or RoseTTAFold ^15^ assesses structural fidelity and stability for early screening. Beyond these efforts, latest methods like ProteinGenerator ^16^, Protpardelle ^17^, BoltzGen ^18^, and RFdiffusion3 ^19^ attempt to jointly design protein sequence and all-atom structure, while cross-modal architectures like ProteinDT ^20^ bridge functional semantics (e.g., “This protein has a lot of *α* -helices”) with protein sequence for natural language-guided sequence generation.

Protein language models (PLM) like ESM2 ^21^, MAGE ^22^, ProTrek ^23^, and ProGen ^24^, as another parallel paradigm, capture evolutionary grammar and residue compatibility through large-scale self-supervised pretraining, enabling high-throughput novel design. To the best of our knowledge, ESM3 ^25^ proposes the first large-scale multimodal PLM (MPLM) that unifies sequence, structural, and functional information within a unified neural network architecture (The largest version has 98 billion parameters). The model processes diverse biological features, including primary sequences, tertiary structures, secondary structure (SS), solvent accessibility (SA), and InterPro annotations ^26^ through specialized tokenization strategies, subsequently constructing a multi-track Transformer framework capable of joint input processing and multimodal output generation. Through self-supervised learning on extensive protein datasets, ESM3 effectively captures cross-modal dependencies and coupling relationships, enabling coherent multimodal protein editing and generation. Consequently, the model successfully explored the latent evolutionary space of protein sequences and generated a functional green fluorescent protein variant with a sequence similarity of 58% to natural counterparts by prompting the core catalytic residues, effectively traversing millions of evolutionary years.

ESM3 has made revolutionary advances in the multimodal editing of general proteins. However, most biological processes involve intricate molecular interactions rather than isolated protein functions. Critical biological systems such as the Cas9-sgRNA complex mediating DNA binding and cleavage through protospacer adjacent moti recognition, hemoglobin’s cooperative oxygen transport via heme-globin coordination, and MHC-TCR-peptide interactions driving immune activation ^27^. All of these crucial activities require precise biomolecular interplay. Designing proteins to interact with various biomolecules is critical for many life science fields, such as drug development and gene editing. The universal ESM3, built on indiscriminate training data, remains underexplored regarding the complex binding modes of different ligands. Moreover, biomolecular interactions are typically guided by conserved motif. Generating novel proteins while understanding these key sites can preserve the native core functions of the template protein and efficiently explore the potential sequence space.

In this study, we propose EiRA, an MPLM specialized for biomolecule-binding protein generation. EiRA undergoes two-stage post-training on ESM3-small (ESM3s, 1.4B): biomolecular-binding domain adaptive masking training followed by binding site-guided preference optimization (named EiRAD). Unconditional generation evaluations reveal that designed proteins exhibit high diversity, novelty, foldability, and functionality. Systematic and multidimensional assessments across 8 test sets demonstrate that our method designs molecular-binding proteins with higher confidence than that of ESM3 (including small/medium/large versions) and RFdiffusion+ProteinMPNN in both single-sequence and complex evaluations. Simultaneously, EiRA’s embeddings effectively characterize natural proteins to support diverse biomolecule-binding downstream tasks, achieving dual benefits in representation and generation. Notably, we identified severe duplicate generation under binding-motif conditions in ESM3 medium (ESM3m, 7B) and large (ESM3l, 98B) models, substantially undermining structural confidence and sequence foldability. This issue emerged during both training stages and was mitigated through repetition penalties and loss adjustment based on direct preference optimization (DPO). Furthermore, whereas ESM3 lacks non-protein biomolecular knowledge, EiRA integrates DNA sequence information via cross-attention to enable multimodal protein editing conditioned on target DNA sequence, addressing critical real-world target-based protein generation scenarios.

To rigorously validate the manufacturability and functional potential of the sequences generated by EiRA, we conducted comprehensive wet-lab characterizations across a diverse set of targets. We challenged the model to explore deep sequence space by designing variants for the RNA-guided endonuclease TnpB and 10 distinct DNA-binding proteins, imposing high mutation rates ranging from 41.82% to 77.30%. Remarkably, experimental validation yielded a 100% success rate in the expression (20/20) and purification (10/10) of these highly divergent variants. In the specific case of TnpB, the generated sequences not only retained expressibility but, in 6 instances, surpassed the expression levels of the wild-type reference. To bridge the gap between static design and dynamic function, we performed all-atom molecular dynamics (MD) simulations on 10 DBP variants. These simulations confirmed that the designed complexes maintain robust interfacial stability and persistent hydrogen-bonding networks throughout 100 ns trajectories, exhibiting conformational resilience comparable to native counterparts. Furthermore, demonstrating EiRA’s generalization capability in *de novo* interaction design, we achieved the “one-shot” generation of a binder for the Glucagon (GCG) peptide. Despite sharing less than 50% sequence identity with its template, the designed binder exhibited a high-confidence structural fold and verified micromolar affinity (*K*_*D*_ = 23.08 *uM*(by Surface Plasmon Resonance (SPR). These results collectively highlight EiRA’s capacity to decouple structural integrity from sequence homology, delivering robust, manufacturable, and functional protein designs without iterative directed evolution.

## Methods

### Data curation

We systematically constructed all datasets for this study through a multi-stage curation pipeline, comprising two training sets for self-supervised adaptation and DPO training and 8 independent biomolecular-binding test sets (see Tables 1 and 2).

**Table 1.** The composition of the pre-training and DPO training sets.

| Data set | UniBind40 (for post-training) | BioDPO (for DPO) |  |  |  |  |  |
| --- | --- | --- | --- | --- | --- | --- | --- |
| Type | Mixed biomolecular binding | DBP | RBP | PBP | ReBP | MBP | PPI |
| $N_{seq}$ | 3,735,303 | 1,214 | 2,332 | 2,420 | 2,319 | 1,419 | 1,047 |
| $N_{res}$ | 1,264,943,457 | 395,728 | 695,230 | 739,251 | 818,405 | 530,400 | 513,642 |
| Mean length | 338 | 326 | 298 | 305 | 353 | 373 | 490 |
| Source | UniProt/AF DB/InterPro | BioLip2/PDB |  |  |  |  | SwissProt/AF DB |

**Table 2.**
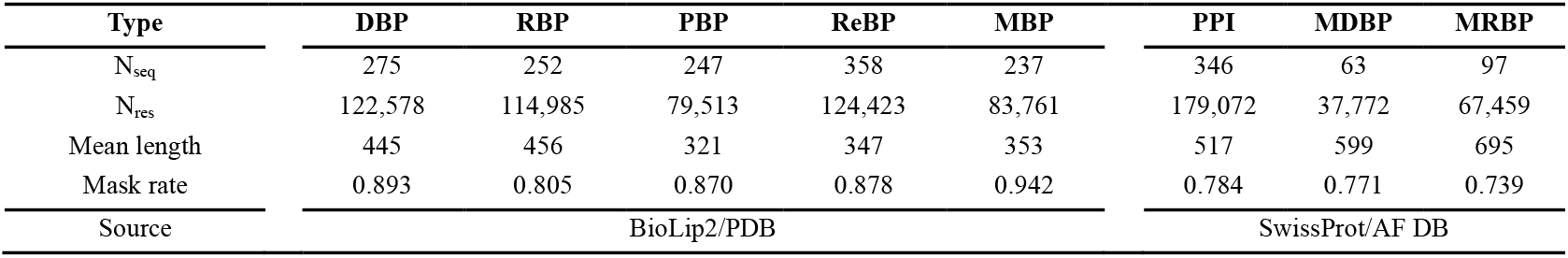
The composition of the 8 test sets across 6 biomolecular types.

| Type | DBP | RBP | PBP | ReBP | MBP | PPI | MDBP | MRBP |
| --- | --- | --- | --- | --- | --- | --- | --- | --- |
| $N_{seq}$ | 275 | 252 | 247 | 358 | 237 | 346 | 63 | 97 |
| $N_{res}$ | 122,578 | 114,985 | 79,513 | 124,423 | 83,761 | 179,072 | 37,772 | 67,459 |
| Mean length | 445 | 456 | 321 | 347 | 353 | 517 | 599 | 695 |
| Mask rate | 0.893 | 0.805 | 0.870 | 0.878 | 0.942 | 0.784 | 0.771 | 0.739 |
| Source | BioLip2/PDB |  |  |  |  | SwissProt/AF DB |  |  |

To align the native knowledge space of the ESM3 with universal biomolecular-binding characteristics, we developed a specialized post-training dataset named UniBind40 via the following protocol (top of Fig. 1): initial ∼54 million biomolecular-interacting protein sequences were retrieved from UniProtKB database ^1^ up to August 17, 2024, followed by MMseqs2 ^28^ clustering at 0.4 sequence identity threshold to derive ∼6.4 million non-redundant cluster representatives. Structural validation proceeded through two pathways: for cluster representatives with existing AlphaFold2 ^29^ predictions (∼4.6 million), we retained only high-confidence structures (pLDDT > 0.7); for sequences lacking precomputed AlphaFold2 structures, we implemented a cascaded prediction strategy using ESM3s for initial high-throughput modeling, followed by ESMFold refinement for cases with suboptimal confidence (ESM3s pLDDT < 0.7), ultimately requiring ESMFold pLDDT > 0.7 for structural acceptance. In addition, samples in which the number of unknown amino acids exceeded 10% of the total sequence length were discarded. This rigorous filtration yielded 3,735,303 qualified biomolecule-binding proteins with 1,264,943,457 residues, designated as UniBind40. The dataset construction emphasizes structural reliability through confidence validation while maintaining sequence diversity via aggressive deduplication.

**Fig. 1.**
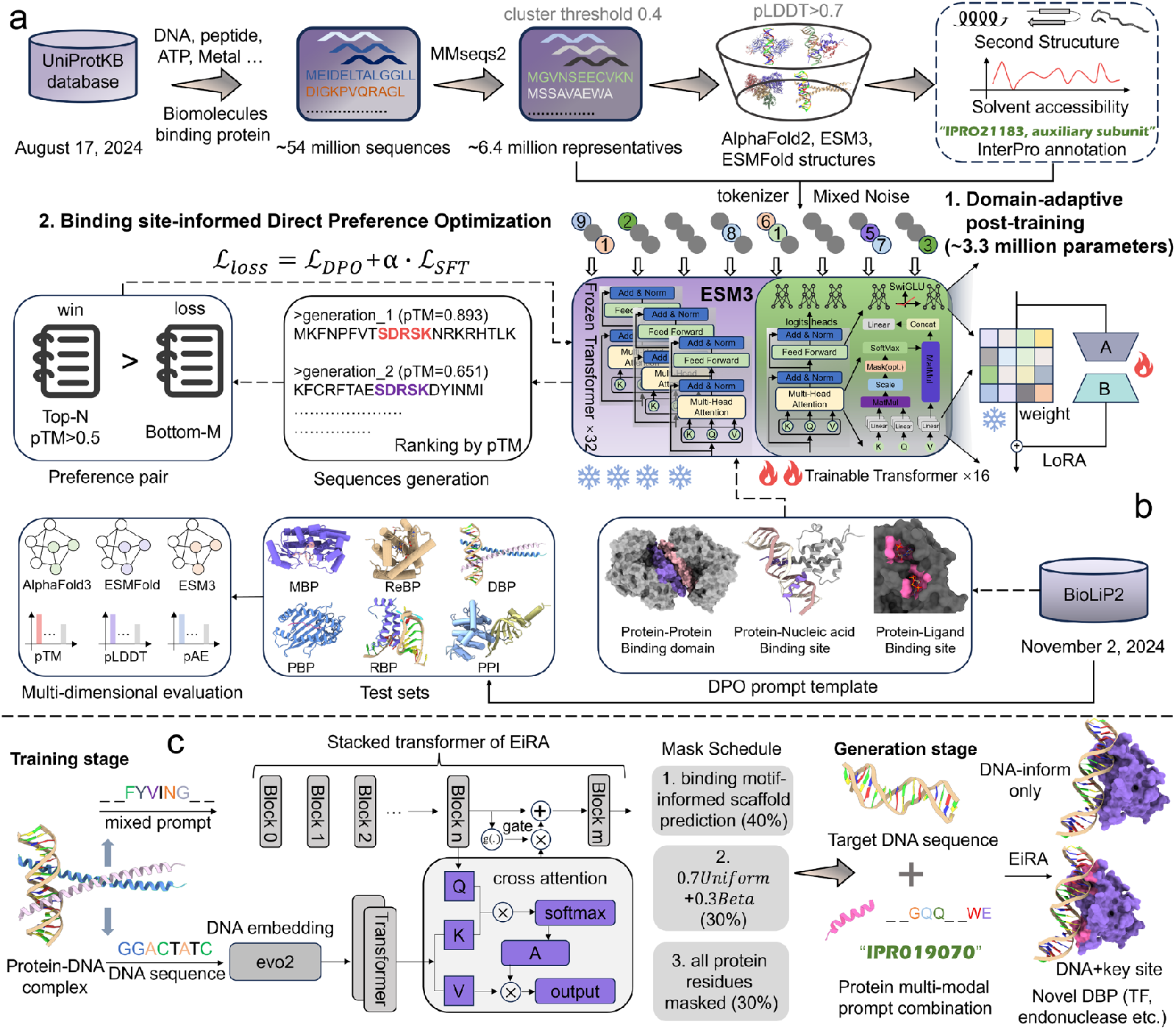
EiRA is a multimodal protein language model specifically customized for biomolecule-binding proteins generation. **a**. We collected ∼54 million universal biomolecule-binding sequences from UniProtKB and selected ∼6.4 million representatives using MMseqs2; then AF2, ESM3, and ESMFold were used to filter proteins with low structural confidence (pLDDT<0.7, remaining 3,735,303, named UniBind40); and finally, corresponding SS, SA, and functional annotations were calculate using the DSSP, Shrake-Rupley, and InterProScan tools. **b**. We first trained the post-16 transformer blocks (LoRA strategy) of ESM3s on UniBind40 by mixing mask noise; then, protein-biomolecule complex data collected from the BioLip database were used to perform a binding site-informed preference optimization. **c**. The large genome language model Evo2 is used to extract the embedding of DNA sequences, which is subsequently introduced into EiRA’s post-4-layer transformer for mask training using a gated cross-attention mechanism. The resulting EiRA model enable DNA-conditioned DBP generation.

Protein-ligand binding data were obtained from the BioLip2 database ^30^ (accessed November 2, 2024), including five categories: DNA-binding proteins (DBPs, 45,686 records), RNA-binding proteins (RBPs, 160,800), Metal-binding proteins (MBPs, 197,122), Peptide-binding proteins (PBPs, 39,065), and Regular-binding proteins (ReBPs, 460,725; e.g., ATP/HEM-binding). After merging redundant chains, we derived 24,459 (DBP), 128,464 (RBP), 125,873 (MBP), 33,755 (PBP), and 252,976 (ReBP) unique chains. Sequence redundancy was reduced using MMseqs2 clustering (40% identity threshold) followed by CD-HIT-2D ^31^ filtering (40% identity) against UniBind40, yielding 1,490 DBP, 2,585 RBP, 1,656 MBP, 2,667 PBP, and 2,677 ReBP chains. Random sampling selected 1,214 DBP, 2,332 RBP, 1,419 MBP, 2,420 PBP, and 2,319 ReBP chains for the DPO training set (named BioDPO), with remaining chains assigned to independent test sets. For protein-protein interaction (PPI) data, sequences with annotated interaction domains were sourced from Swiss-Prot ^32^. After redundancy removal with UniBind40 and filtering for AlphaFold2 pLDDT>0.7, 1,047 sequences were added to BioDPO, and 351 were reserved for the PPI test set.

Independent test samples for DBPs, RBPs, MBPs, PBPs, and ReBPs employed binding motif prompts containing sequences and coordinates of 4 flanking residues around the real binding site (8 for MBPs/ReBPs). PPI test samples directly utilized full domain sequences/coordinates as prompts. To evaluate multi-domain design capability, multi-domain DBP/RBP test sets were constructed by: (1) selecting UniProtKB entries with ≥2 annotated domains; (2) removing redundant sequences and those with max identity>40% against UniBind40 via MMseqs2 and CD-HIT-2D; (3) retaining sequences with total domain length <40% of the full sequence. This produced 97 multi-domain RBPs and 64 multi-domain DBPs, processed with domain-level prompts following PPI samples in BioDPO. All of this data is available at https://github.com/pengsl-lab/EiRA. All feature generation protocols can be found at Supplementary Text S1.

### Evaluation metrics

For a designed protein sequence, we used three models, i.e., **ESM3 structure decoder, ESMFold** ^21^, and **AlphaFold3** (AF3, utilizing Boltz-1, an open-source re-implementation of the AF3 ^33^) to predict its tertiary structure and obtained a series of metrics. The ESM3 structure decoder is used to convert the structural tokens output by MPLM into backbone atomic coordinates; ESMFold is an MSA-free structure predictor based on a protein sequence language model; AF3 is a diffusion model-based MSA-depend complex structure predictor. The evaluation of the designed sequences by different structural predictors can prevent biased assessments of a single structure prediction model. Specifically, predicted Template Modeling Score (**pTM ↑**) assessed the overall stability and rationality of the predicted structure; predicted Local-Distance Difference Test (**pLDDT ↑**) evaluated the local positional validity of residues; and predicted Aligned Error (**pAE ↓**) depicts the confidence of the relative positions of pairs of residues in the predicted structure. In addition, we calculated the self-consistent Root Mean Square Deviation (**scRMSD ↓**) of the designed backbone against the ESMFold and AF3 structures to assess the designability of the backbone.

For the evaluation of the design sequence and its ligand binding ability, the interface pTM (**ipTM ↑**), interface pLDDT (**ipLDDT ↑**), and interface pAE (**ipAE ↓**) of AF3 were employed to assess the accuracy of the interface interactions between subunits in the predicted complexes. Additionally, we conducted sequence validation using three complementary approaches: CD-HIT clustering to assess **diversity**, MMseqs2 similarity searches to evaluate **novelty**, and InterProScan scans to verify **functionality**.

### Domain-adaptive self-supervised post-training

The general MPLM ESM3 has learned a vast amount of multimodal world knowledge about proteins but lacks focus on specific domains such as biomolecular-binding. To transfer the general ESM3 knowledge space and focus it on the field of biomolecular binding, we continue to perform a mask-based self-supervised post-training strategy for ESM3-small (ESM3s) MPLM on our well-curated UniBind40 training dataset. Although ESM3 includes three versions: ESM3s, ESM3-medium (ESM3m), and ESM3-large (ESM3l) with 1.4, 7, and 98 billion paramters respectively and Hayes *et al*. ^25^ reports that larger model have stronger generative capabilities, only the 1.4B version of ESM3 has open weights and source code.

For different modalities, we employed different noise scales as ESM3 to simulate conditional generation tasks. Unlike ESM3, for the sequence track, we used a betalinear70 noise composed 30% *βeta*(3,9) and 70% *Uniform*(0,1), which offers a higher mask rate than ESM3 training strategy to optimize generation tasks (Fig. S1). We experimented with various fine-tuning strategies, including directly updating the original weights, Low-Rank Adaptation (LoRA) ^34^, and various variants. Ultimately, it was found that the vanilla LoRA achieved the best performance. Previous study has demonstrated that partial parameter fine-tuning can outperform full parameter fine-tuning on specific tasks and datasets ^35^. Here, we apply LoRA to the last 16 transformer blocks and the token classification head of each track, totaling 3,368,046 trainable parameters (only 0.24% of all ESM3s 1.4B parameters). The rank of LoRA is set to 8, the scaling factor *a* is 32, and the dropout rate is 0.2. Cross-entropy is used to calculate the loss between the true tokens and the predicted tokens for each track. Since we observed a severe phenomenon of repeated generation of tokens conditioned at binding sites in ESM3, if the predicted tokens at 7 consecutive adjacent positions are consistent, the loss at these positions is multiplied by a penalty factor of 2. The 8-bit quantized AdamW optimizer is used to update the parameters, with a learning rate of 4e-04, *β*_1_ of 0.9, *β*_2_ of 0.95, and a weight decay of 5e-04. The cropping length is 768. Long sequences are split, and short sequences are filled with <pad> token. The batch size is 160. We implemented training using the standard Distributed Data Parallel strategy of Pytorch and employed a mixed precision training with BF16 and FP32. All training was performed on 8 NVIDIA H20 GPUs.

### Binding site-informed preference optimization

The recent DeepDeek-R1 achieved SOTA performance for LLM using only a RL strategy called Group Relative Policy Optimization without any supervised fine-tuning (SFT) ^36^. This concept has been borrowed in the field of PLM ^25, 37^. ESM3 uses an Iterative Reasoning Preference Optimization ^38^ strategy composed of a Direct Preference Optimization (DPO) ^39^ loss and an NLL loss conjunctions to bias the model towards design proteins with high pTM and low backbone constrained site RMSD (cRMSD). Here, we employ the DPO and SFT strategies to enhance generation quality under biomolecule binding site conditions.

#### Algorithm description

To enhance the comprehensive generation quality of EiRA under the condition of binding motif, we initially used only the DPO strategy to train the EiRA model, but crashed duplicate generation occurred similar to ESM3. Therefore, we introduce an additional SFT loss to impose a larger penalty on mispredictions of repeatedly generated regions. Consequently, we propose a binding site-informed DPO strategy combined with an SFT loss as follows (Equations 1, 2, and 3):

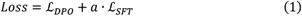

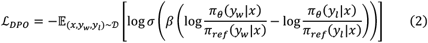

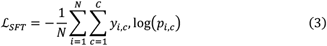

where *π*_*θ*_ and *π*_*ref*_ are respectively the learnable policy model and the frozen reference model; *x, y*_*w*_, and *y*_*l*_ are the input prompt sequence, the chosen and rejected designed sequences separate; *a* is the weight of SFT loss and set to 0.2; *b* is used to control the degree of deviation between the policy model and the reference model and set to 0.1. ℒ_*SFT*_ directly employs a multi-class cross entropy. *N* is the total number of residues in current batch; *C* is the number of categories; *y*_*i,c*_ indicates whether the residue *i* belongs to category *c*, and *p*_*i,c*_ indicates the probability that the residue *i* is predicted to belong to category *c*. To suppress duplication, when the maximum predicted probabilities of 5 consecutive positions (or more) all point to identical amino acid type, the loss of this region is multiplied by a penalty coefficient of 2.

#### Preference pair generation

For BioDPO, prompts were derived from the binding site coordinates and residues, supplemented by *n* flanking positions (*n* = 4 for DBP/RBP/PBP; n=8 for MBP/ReBP), with the remaining regions masked for reconstruction. We generated 30 variants per prompt and constructed preference pairs using a multi-stage filtering protocol. First, valid batches were identified using dynamic pTM thresholds (*p, M, q*): if the maximum pTM> *p*, the top-*M* variants were labeled as “preferred” (provided the lowest pTM> *q*) and the bottom-6 as “dispreferred”. Three parameter configurations were employed: (0.7, 5, 0.6), (0.6, 4, 0.5), and (0.5, 3, 0.5). Second, to ensure data quality, we discarded entire batches where the minimum pTM>0.8 or the maximum pTM<0.5. Finally, sequences were deemed negative if they contained >8% unknown residues or exhibited high repeatability (defined as having a repetitive substring >7 residues and the top-2 amino acids constituting > 40% of the sequence). This pipeline resulted in a final dataset of 224,891 preference pairs.

## Results

### EiRA generates biomolecular binder with higher fold confidence

We assessed the generative capabilities of EiRA against ESM3 using three distinct evaluation strategies: (1) unconditional generation, (2) single-sequence scoring, and (3) evaluation of binding ability to specific biomolecules.

#### Unconditional generation

Unconditional generation is a key benchmark for protein design models, as it evaluates their capacity to navigate the vast potential sequence space. In this task, the model generates sequences based solely on a specified length, with all other contextual information masked. We generated a dataset of 100,000 sequences by first sampling 1,000 distinct lengths from three intervals: [50, 500], [501, 1000], and [1001, 1500], with sampling probabilities of 0.6, 0.3, and 0.1, respectively. For each sampled length, 100 unique sequences were generated. Subsequently, the tertiary structure for each of these 100,000 designed sequences was predicted using the ESM3 structure decoder to calculate key quality metrics (see Fig. 2a). EiRA demonstrated superior performance, achieving average pTM and pLDDT scores of 0.473 and 0.707, which are 35.70 and 65.38% higher than those of ESM3s, respectively. A direct comparison of metrics for sequences of identical lengths reveals that EiRA consistently outperforms ESM3 across all length ranges (see Fig. 2d). An observed trend for both models is the degradation of pTM and pLDDT scores with increasing sequence length, suggesting that the modeling of long-range interactions remains a challenge for both generative and predictive models.

**Fig. 2.**
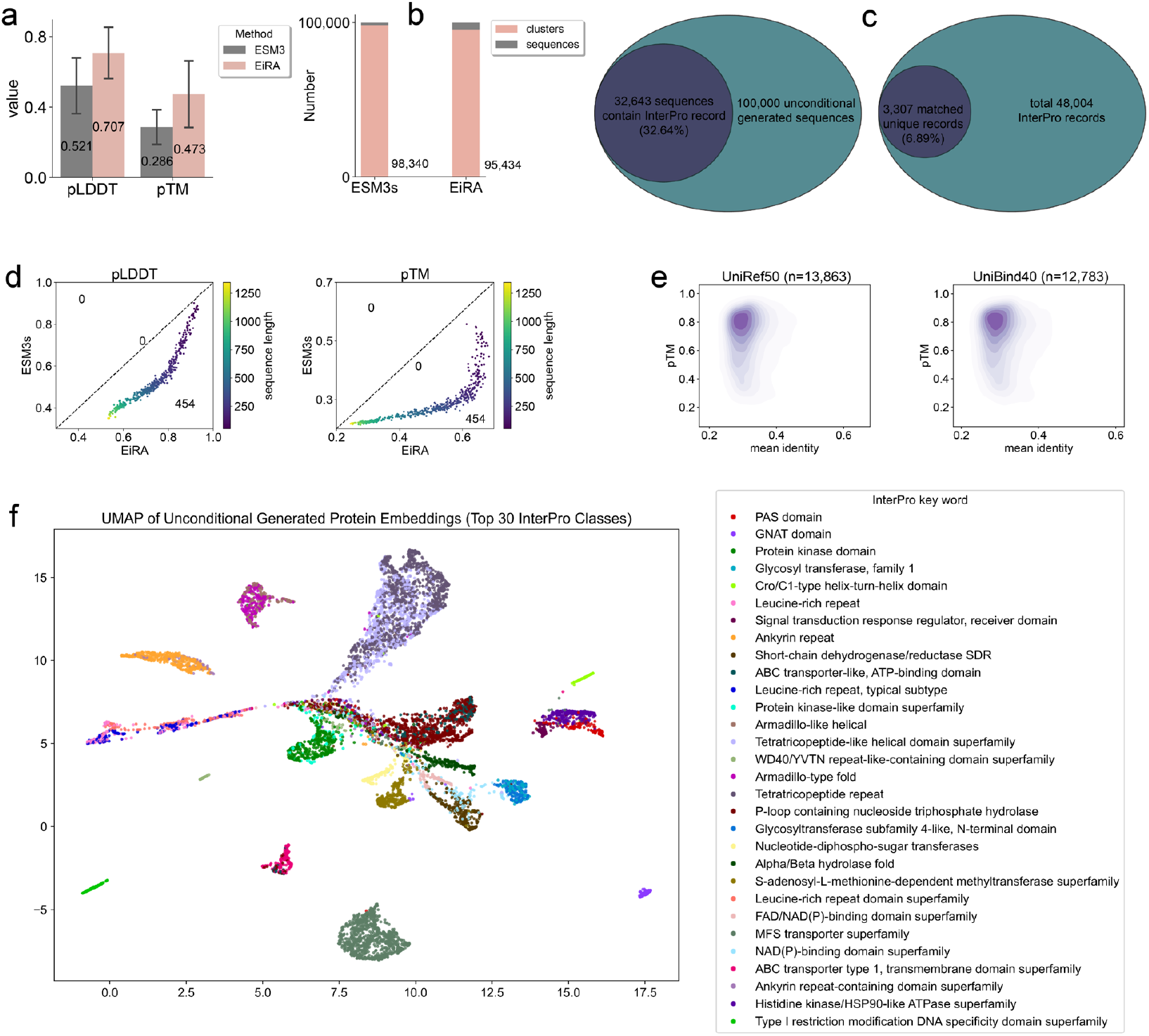
Unconditional generation evaluation. **a**. Direct comparison of pLDD and pTM of 10,000 unconditional generated sequences. **b**. Number of clusters of 100,000 generated sequences (CD-HIT tool with a sequence identity threshold of 0.4 for clustering). **c**. Venn diagram of the InterPro annotation of the designed sequence. **d**. Head-to-Head comparison of mean pTM (or pLDDT) for sequences of equal length. **e**. Kernel density for the mean sequence identity (against UniRef50 and UniBind40 respectively, using MMseqs2 easy-search) and pTM. **f**. Embedding visualization of sequences with top-30 InterPro annotation numbers, including several typical biomolecular-binding domains.

The **diversity** of generated sequences is a critical indicator of a model’s generative scope. Following the methodology in Proteína ^8^, we quantified diversity by clustering the 100,000 designed sequences using the CD-HIT program with a sequence identity threshold of 40%. This analysis yielded 95,434 clusters for EiRA and 98,340 for ESM3s (see Fig. 2b). A high cluster count signifies low sequence similarity among the generated samples. In addition, the MMseqs2 tool was used to search the similar sequences of 100,000 designed sequences against the UniRef50 and UniBind40 databases. Consequently, for UniRef50 and UniBind40, 13,863 and 12,783 designed sequences that were matched to a total of 2,933,813 and 1,495,174 records, respectively. We averaged the identities of the different records matched to the identical sequence and plotted the kernel densities against pTM (see Fig. 2e). The densities in the upper left corner indicate generally **low similarity** and **high structural confidence**. These results indicate that EiRA not only designs structurally superior proteins but also maintains a high degree of sequence diversity and novelty, comparable to that of ESM3s.

To validate **functionality** across 100,000 unconditionally generated sequences, we performed functional annotation prediction using InterProScan. Of these, 32,643 sequences matched known InterPro entries (see left of Fig. 2c), yielding 149,571 redundant InterPro records (Fig. S7) comprising 3,307 unique IDs (see right of Fig. 2c). This demonstrates EiRA’s capacity to generate functional protein sequences unconditionally. We ranked the top-30 non-redundant InterPro annotations (assigning each protein its most frequent InterPro ID) and visualized corresponding sequences via UMAP (see Fig. 2f). Notably, these include multiple biomolecule-binding domains (PAS, ATP-binding, and Cro/C1-type HTH domains), exhibiting distinct clustering patterns. The PAS domain (ubiquitous in eukaryotes and prokaryotes) mediates protein homo-/heterodimerization, while the Cro/C1-type HTH domain represents one of nature’s most efficient compact DNA recognition modules. Visualization of 16 representative samples (Fig. S6) further confirms EiRA’s generation of biomolecule-binding domains (targeting DNA, RNA, ATP, GTP, UDP, NAD, and AMP), including classical DNA-binding architectures (HTH, BTB, and zinc finger C2H2 domains) with rich secondary structure. Significantly, gen_545045 (Fig. S6, lower right) integrates three distinct domains (NAD-binding, UDP-binding, and a central domain), highlighting EiRA’s capacity for complex multi-domain protein generation.

Collectively, large-scale validation demonstrates that EiRA unconditionally generates proteins with enhanced structural confidence and biomolecule-binding functionality over ESM3s, while maintaining high diversity.

#### Monomer evaluation

This experimental protocol involved systematic evaluation of five MPLMs across 8 benchmark datasets: ESM3 small (1.4B, ESM3s), ESM3 medium (7B, ESM3m), ESM3 large (98B, ESM3l), EiRA without DPO (1.4B), and DPO-enhanced EiRA (1.4B, EiRAD). In addition, a mainstream scheme, i.e., RFdiffusion (backbone generation)+ProteinMPNN (sequence design), was used as the SOTA control. Generated sequences were subsequently processed through three structural predictors to obtain quantitative metrics (Fig. 3a).

**Fig. 3.**
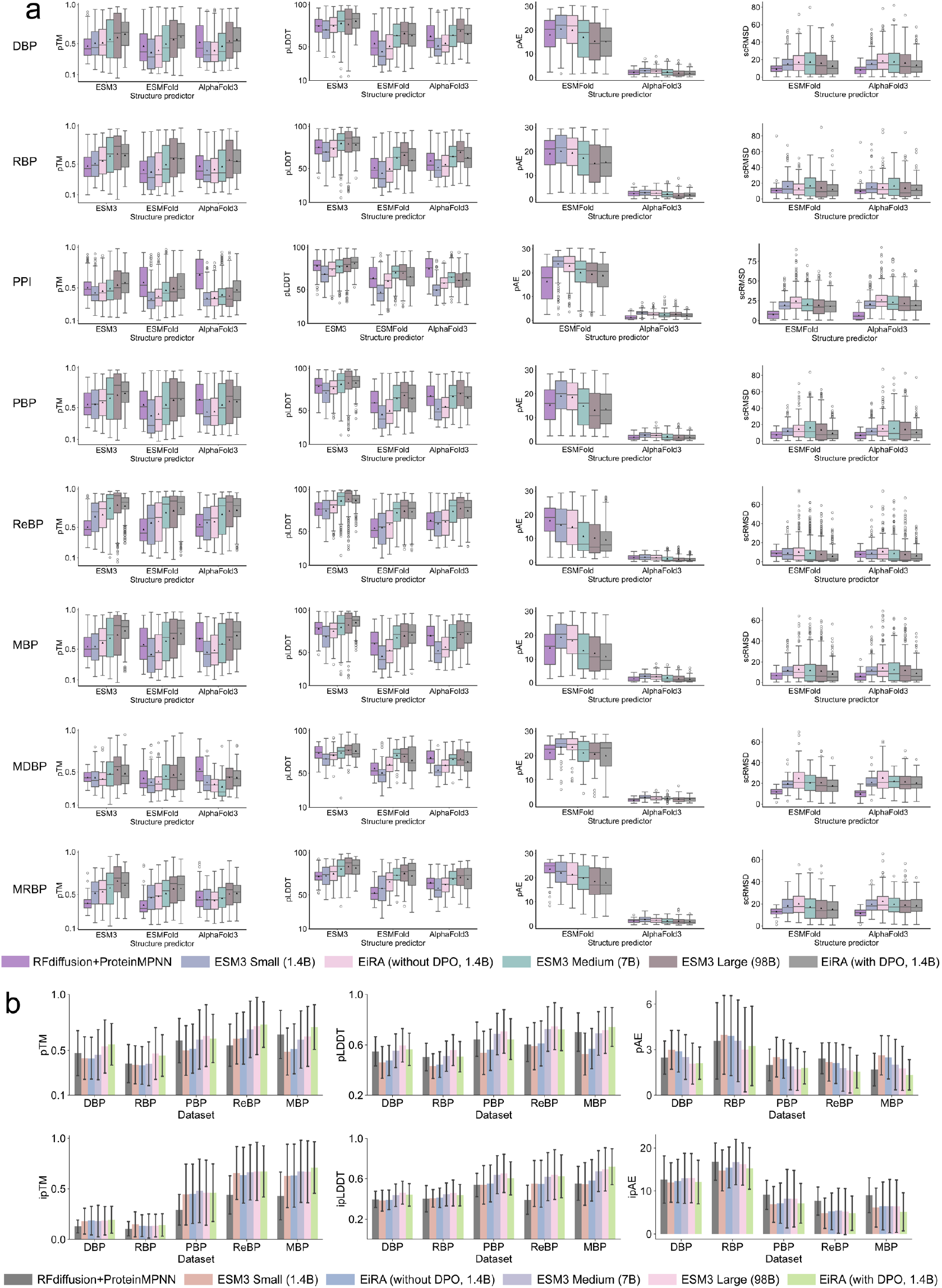
EiRA outperforms ESM3 on biomolecule binder generation over single sequence (a) and complex evaluation (b). Since ESM3 medium and ESM3 large are not open-source, the generations come from the Forge service at https://forge.evolutionaryscale.ai/. Boltz was employed as a replacement for AF3. MSA for complex structure prediction was generated using MMseqs2 against UniRef50 and Small BFD (taking only the first non-consensus sequence from each cluster in BFD) database.

Notably, EiRA without DPO demonstrated superior performance over ESM3s, confirming that domain-adaptive post-training effectively transfers MPLM knowledge to biomolecular binding contexts, yielding more physiologically plausible biomolecule-binding protein designs. The observed performance hierarchy (ESM3l >ESM3m > EiRA >ESM3s) aligns with established scaling laws ^40^, where increased model parameters correlate with enhanced learning capacity. However, EiRAD produced sequences with metric improvements surpassing ESM3m and achieving parity with ESM3l. Intriguingly, comparative analysis revealed EiRAD’s superior pTM scores versus ESM3l (indicative of enhanced global stability) alongside lower pLDDT values (suggesting reduced local structural precision). Both ESMFold and AF3 evaluations confirmed EiRAD’s design superiority through significantly lower scRMSD values, validating methodological reliability.

Dataset stratification revealed pronounced metric disparities: MBP and ReBP ligands exhibited elevated pTM/pLDDT values, likely attributable to evolutionary conservation facilitating pattern recognition in training data, and the average length is relatively short. Conversely, PPI, MDBP, and MRBP designs showed reduced performance, potentially stemming from template sourcing limitations (AlphaFold DB vs. experimental PDB structures) that propagate structural inaccuracies. Crucially, EiRAD achieved ESM3l-comparable pTM scores in multi-domain protein tests, demonstrating effective handling of complex structural architectures.

An analysis of performance reveals a notable exception to EiRAD’s general superiority: it is inferior to RFdiffusion on the PPI and MDBP test sets. We attribute this to the nature of these two benchmarks, which contain a relatively large number of prompt residues (prompt rates: 21.59 and 22.89%) that heavily constrain the design task. RFdiffusion’s PDB-centric training predisposes it to generate sequences that conservatively “infill” the structural scaffold in a manner similar to natural proteins. This yields designs that are readily validated with high confidence by structure predictors. Conversely, EiRA, which leverages a more diverse set of predicted structures, maintains its tendency to generate novel sequences. For these specific, highly-constrained tasks, this novelty is a double-edged sword. The resulting unfamiliar sequences challenge structure predictors, leading to lower confidence scores and masking EiRA’s underlying generative capability.

In general, following domain-adapted post-training, the EiRA framework exhibited enhanced capabilities in biomolecular-binding protein design compared to the native ESM3s architecture. Building upon this foundation, the binding site-informed DPO-enhanced EiRAD demonstrates remarkable parameter efficiency, achieving performance surpassing (or parity with) the 98B-parameter ESM3l model despite operating with merely 1.4B parameters. Additionally, EiRAD’s superior performance over the SOTA method, RFdiffusion, across most test sets underscores the effectiveness of our method.

#### Complex Evaluation

We input the designed sequences from the DBP, RBP, PBP, ReBP, and MBP test sets, along with their respective ligand chains in the PDB template, into AF3 to obtain predicted complex structures and evaluation metrics (see Fig. 3b). EiRA consistently outperformed the original ESM3s models across all test sets in terms of pLDDT, even without DPO enhancement. Furthermore, the DPO-enhanced EiRAD surpassed ESM3m and ESM3l in multiple comparisons. For instance, EiRA achieved higher pTM than both ESM3l and ESM3m on three out of five test sets (DBP, ReBP, MBP). Additionally, EiRA with DPO exhibited lower ipAE than ESM3m and ESM3l across all test sets. Notably, EiRA demonstrated superior performance on the MBP test set for all metrics, indicating unique advantages. Regarding ipTM, while EiRA was significantly higher than other methods on MBP, performance differences were negligible on other datasets. This pattern may arise since the generative prompts directly inherit interface residue properties from the PDB templates. We also observed performance that was comprehensively superior to RFdiffusion, both in terms of overall pTM and interface ipTM scores.

We visualized four designed cases: an RBP (PDB ID: 6dt8, Bacteriophage N4 RNA polymerase II elongation complex 1), a DBP (PDB ID: 1gm5, RecG protein, DNA-specific helicase), an MBP (PDB ID: 8on7, FMRFa-bound Malacoceros FaNaC1), and a PBP (PDB ID: 6wtb, Sort-Tagged Drosophila Cryptochrome). As shown in Fig. S2-S5, EiRA generated proteins with higher confidence (pTM: 0.79, 0.73, 0.828, 0.901; ipTM: 0.84, 0.68, 0.912, 0.835) than all ESM3 versions in all cases. Crucially, AF3 predicted successful target ligand binding for all four EiRA designs, whereas designs for the RBP (by ESM3l) and DBP (by ESM3m and ESM3s) were predicted not to bind their target nucleic acids. For DBP (1gm5) and MBP (8on7), the structural confidence of the EiRA generations is even higher than that of the AF3 structure of the natural sequence. Furthermore, EiRA designs maintained contact between the key prompt motif (highlighted in pink) and target ligand in the RBP, PBP, and MBP cases, indicating successful inheritance of the template’s core interactions.

Low sequence identities (18.37, 17.73, and 19.60%) for these three cases confirm the ability to explore broader sequence and structural space while preserving original interaction.

### Relief from repeated generation

Repetitive generation in LLMs has long been recognized within NLP. In our experiments, significant repetitive generation was observed in biomolecule-binding motif-conditioned ESM3, particularly in the medium (ESM3m) and large (ESM3l) versions. This resulted in a severe skew in amino acid type distribution (Fig. 4a), with ESM3l and ESM3m exhibiting a pronounced bias towards Alanine and Leucine, respectively. In contrast, ESM3s and EiRAD produced more balanced distributions. Such bias towards specific residues compromises sequence diversity and foldability. Formally, a sequence was defined as repetitive if its longest single-amino acid repeat substring exceeded length 5 and the combined frequency of the top two amino acids exceeded 40% of the protein length. Counting repetitive sequences generated by different models across 8 test sets (Fig. 4b) revealed three key phenomena: **1**. ESM3l and ESM3m generated a high number of repetitive sequences (742 and 796, respectively); **2**. post-training directly on ESM3s exacerbated repetitive generation; and **3**. our proposed loss-penalization and DPO+SFT strategies effectively mitigated repetitive generation during post-training and DPO. Sequences designed by ESM3l and ESM3m were categorized into repetitive and non-repetitive groups. Visualization of their pLDDT and pTM scores (Fig. 4c) shows that the repetitive group scored significantly lower on both metrics than the non-repetitive group, indicating reduced structural confidence in repetitively sequences.

**Fig. 4.**
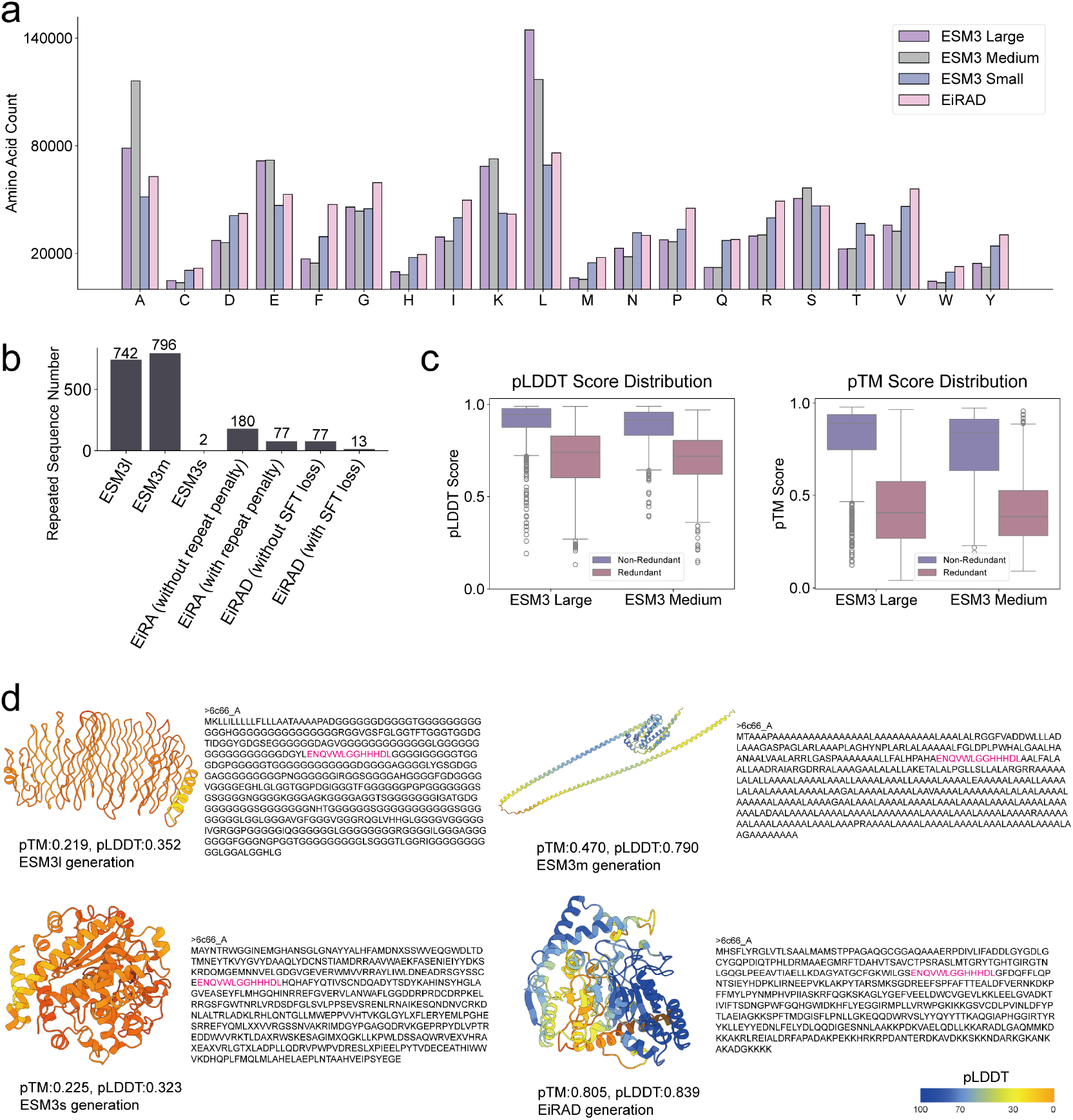
EiRAD rescues the duplicate generation. **a**. Amino acid distribution of the generated sequences for the test sets. **b**. Statistics on the number of duplicate sequences for the test sets. **c**. Repeated designed sequences lead to extremely deteriorated structure confidence. **d**. The case (RNA-binding protein, PDB ID: 6c66_A) of repeated sequences generated by ESM3l and ESM3m. The prompt residues are highlighted in pink. The structures are predicted by Boltz.

Illustratively, the structures generated for target 6c66_A (chain A of CRISPR RNA-guided surveillance complex) by the four models clearly demonstrate ESM3’s failure modes (Fig. 4d). Under identical RNA-binding residue prompts, ESM3l and ESM3m exhibited biases towards Glycine and Alanine, respectively, failing to form a stable tertiary structure. Although ESM3s avoided repetitive generation, their structural confidence remained low (pLDDT: 0.323, pTM: 0.225). Conversely, EiRAD generated proteins with diverse amino acid residues that formed a stable tertiary structure (pLDDT: 0.839, pTM: 0.805). In conclusion, the higher-parameter ESM3l and ESM3m models, typically considered more generative than ESM3s, suffer from severe repetitive generation under biomolecule-binding motif conditioning. Our improved loss strategy mitigates the repetitive generation arising during ESM3s-based training, achieving simultaneous improvements in structural confidence and amino acid diversity.

### Ablation study on fine-tuning strategy and parameter

ESM3 is trained on multiple databases, including UniRef ^41^, Mgnify ^42^, JGI ^43^, OAS ^44^, and PDB ^45^, with approximately 3,143 million unique samples. Since our training set UniBind40 is only a specific subset of the natural protein space, directly fine-tuning all parameters may compromise the fundamental biochemical and protein knowledge held by the ESM3. Here, we observe the performance of the generation task with different levels of fine-tuning parameter counts (see Table S1). Most intuitively, the improved structural confidence compared to ESM3 after fine-tuning demonstrates the advantages of our domain-adaptive training strategy. Furthermore, fine-tuning 16 (or 32) Transformer blocks achieves leading structural confidence. Considering that the latter requires more training and inference resources, the former is easier to scale, optimize, and apply in practice.

In addition, we performed different fine-tuning schemes and observed the generated performance (see Table S2). It is worth noting that directly updating the original ESM3 weights resulted in mediocre performance. Among the many LoRA techniques (vanilla LoRA ^34^, LoRA+ ^46^, LoKr ^47^, hydraLoRA ^48^, and AdaLoRA ^49^), vanilla LoRA demonstrates higher structural confidence than other variants. A potential reason may be that these variants are task-specific and lack generalizability.

### Representation learning for downstream prediction task

PLMs encode evolutionary information and biochemical features within protein families, and their embedding representations support diverse downstream tasks, such as secondary structure prediction, tertiary structure prediction, and mutation effect prediction. ESM3 demonstrated that high mask rates improve generation performance but impair representation learning. In the EiRA training strategy, we therefore increased the mask rate to optimize the generative task. Here, we evaluate EiRA’s representation learning capability specifically for biomolecular binding prediction tasks (see Fig. 5). We extracted a feature matrix of size *L* × 1536 from the final transformer block of both ESM3 and EiRA. The normalized matrix was then applied to four downstream tasks: DNA-binding interface (DBI), RNA-binding interface (RBI), ATP-binding interface (ABI), and DNA-binding protein (DBP) prediction. Details of the training and test sets are provided in Table 3. To accommodate different scenarios: for DBI and RBI prediction, we employed an EGNN model based on tertiary structure; for ABI prediction, a lightweight sequence-based BiLSTM model was used; and for the DBP task, we first averaged the feature matrix along the first dimension (sequence length) before feeding it into a simple multilayer perceptron (MLP) model. EiRA consistently outperformed ESM3 across most metrics and datasets. For instance, EiRA achieved AUPR values of 0.577 (DNA129), 0.387 (DNA181), 0.386 (RNA117), 0.730 (ATP41), and 0.656 (ATP202), representing improvements of 3.40, 2.65, 6.33, 2.24, and 1.39%, respectively, over ESM3 (Fig. 5a). Furthermore, for DBP prediction on UniSwiss-Tst, EiRA provided higher prediction probabilities for 196 DBPs and lower probabilities for 286 non-DBPs (Fig. 5e).

**Table 3.**
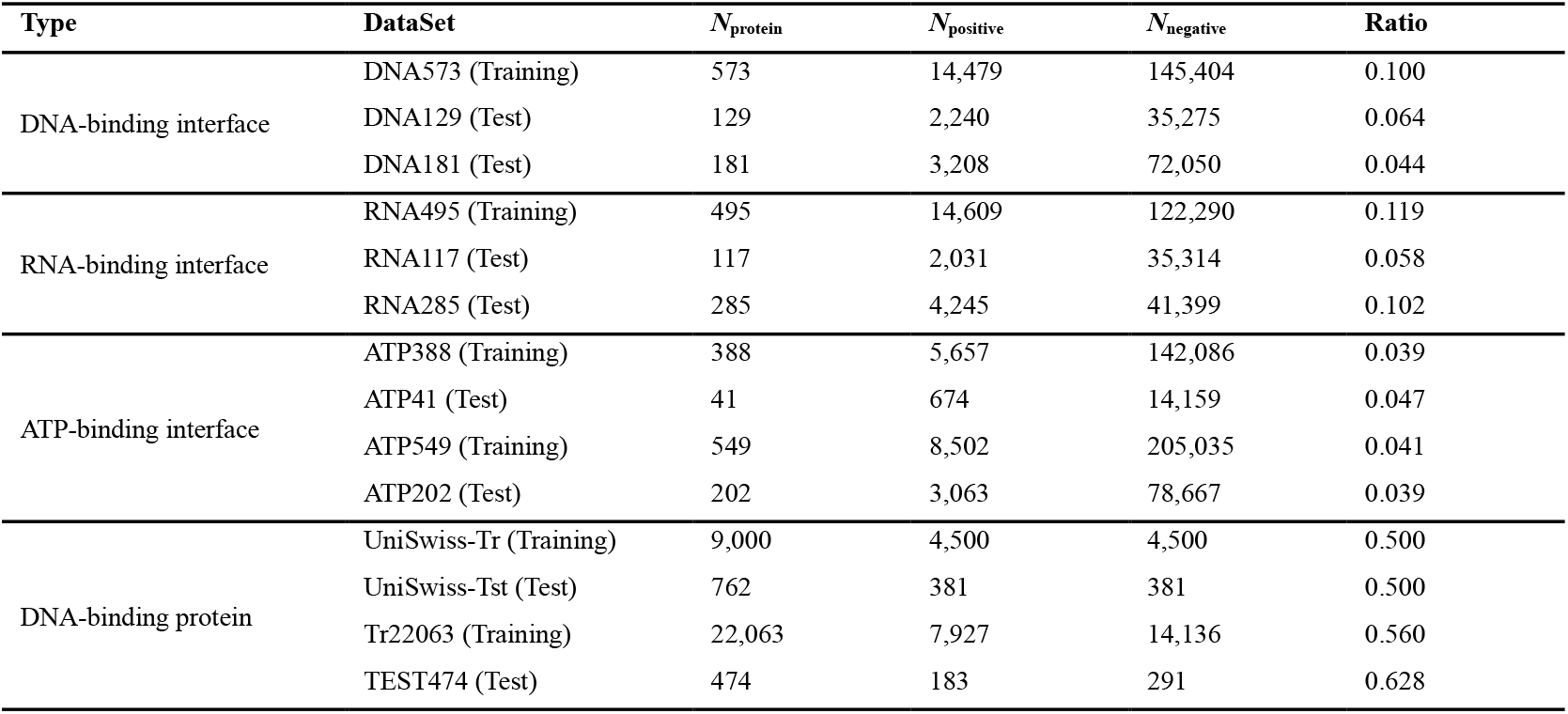
The composition of the datasets for representation learning.

**Fig. 5.**
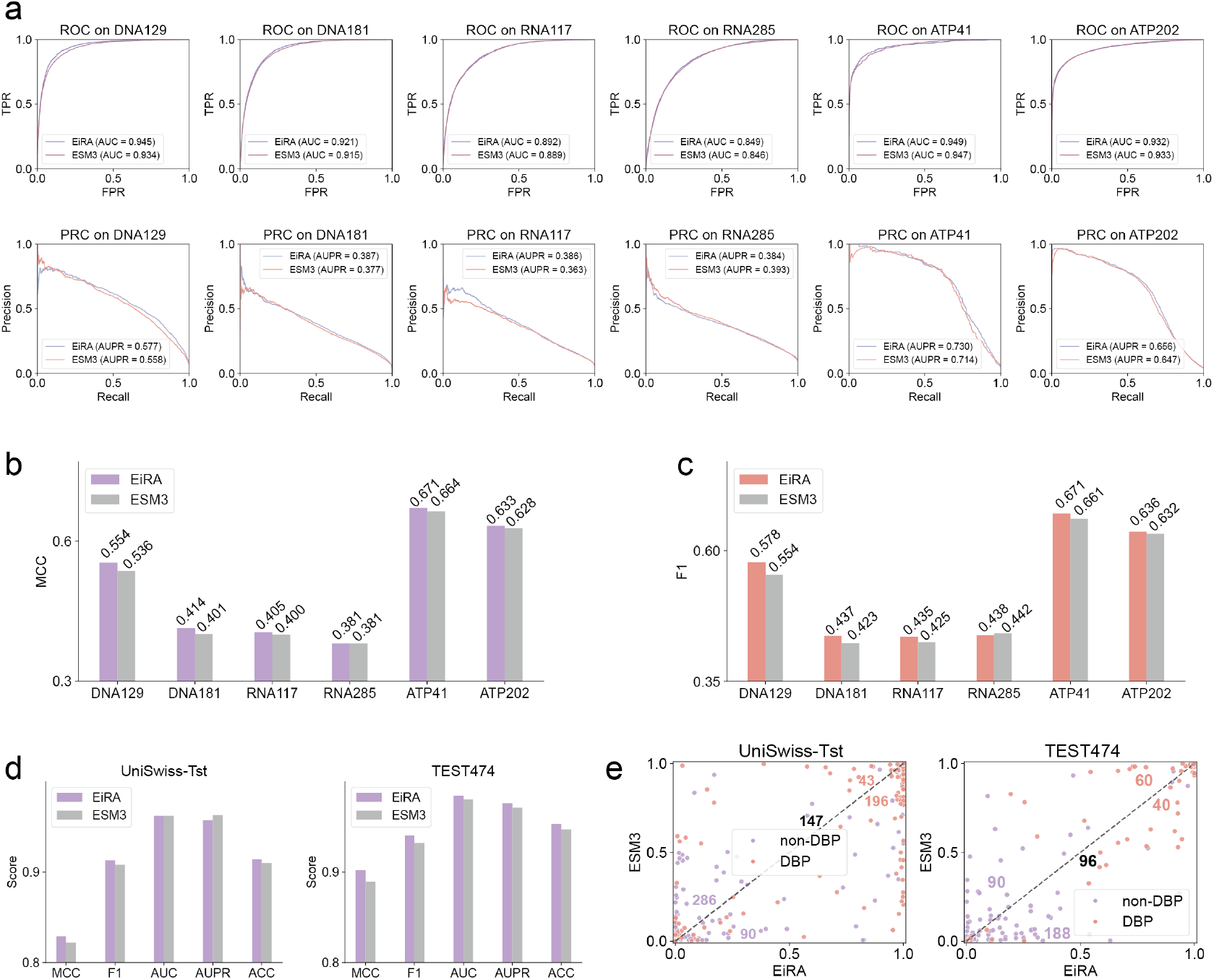
Representation learning evaluation on four biomolecules-binding-related downstream tasks. **a-c**. Evaluation on 6 test sets for DNA-binding, RNA-binding, and ATP-binding interface predictions. **d**. Evaluation on 2 DNA-binding protein prediction test sets, i.e., UniSwiss-Tst and TEST474. **e**. Head-to-head comparison of the prediction probabilities between EiRA and ESM3 embedding on 2 DBP prediction test sets. Each point represents a protein, and the number indicates the number of DBP or non-DBP located in the diagonal, upper, or lower triangle.

Furthermore, we present three representative cases (Fig. S8): the HSDR subunit of the ECOR124I restriction enzyme (ATP-binding, PDB ID: 4XJX_B), the DNA repair protein RAD4 (DNA-binding, PDB ID: 7m2u_A), and Mitochondrial ribosomal protein L45 (12s rRNA-binding, PDB ID: 6ydw_Bi). EiRA achieved MCC values of 0.529, 0.592, and 0.471 for these samples, respectively, exceeding those of ESM3 (0.280, 0.202, and 0.000). Specifically, ESM3’s high number of false negatives (FNs) on 7m2u_A indicates frequent misclassification of non-DNA-binding sites (non-DBS) as DNA-binding sites (DBS), a failure largely avoided by EiRA. For 6ydw_Bi, a component of the large mitochondrial ribosomal complex, ESM3 failed to identify any RNA-binding residues. Conversely, EiRA successfully predicted multiple native binding sites; moreover, its predicted FN residues clustered near the target rRNA, indicating its distinct advantages indicating the distinctive advantage. Collectively, the above experiments demonstrate that EiRA, enhanced by training on numerous molecular binding proteins, captures meaningful biological patterns that safeguard representation learning capability while simultaneously enhancing generative task performance.

### DNA sequence-guide binder generation

Target-based binder generation represents a core challenge in protein design ^2-4^. While ESM3 leverages extensive protein data, it lacks information on other biomolecules. Here, take DNA as an example, we integrate DNA embedding knowledge from Evo2 ^50^ (a SOTA DNA language model) into ESM3 as a generative condition via cross-attention (Fig. 1c). Specifically, Evo2 embeddings are first processed through a 2-layer transformer block. The outputs of EiRA’s final four transformer blocks are then residually fused with the DNA transformer via gated cross-attention, and the combined representation is used for masked pre-training. Three distinct masking strategies were applied: (1) Masking residues at untied localization sites (40% of cases); (2) EiRA’s original pre-training masking strategy (30% of cases); (3) Full masking of protein residues with DNA input retained for DNA-only generation (30% of cases). We fine-tuned the final 32 transformer blocks of EiRA using all 1,214 DNA-binding proteins (DBPs) from BioDPO. DNA sequences exceeding 50 residues (the average training length is 26) were truncated; shorter sequences were zero-padded. Protein processing mirrored EiRA’s original protocol.

Using binding sites and DNA chains from BioDPO complexes as prompts, we generated 101,950 sequences (50 per prompt). MMseqs2 searches against UniRef50 and UniBind40 identified similar sequences for 83,280 and 75,725 samples, respectively. Kernel density plots of mean identity versus pTM reveal a pronounced cluster in the upper-left quadrant, indicating designed sequences achieve both novelty and high predicted structural confidence (Fig. 6b). UMAP visualization further compares generated sequences against EiRA embeddings of UniDBP40 (from ESM-DBP ^35^. Clustering all UniProt DBPs at 40% similarity (CD-HIT) yielded 170,264 representatives native DBPs.), assessed by sequence identity (Fig. 6c) and pTM (Fig. 6d). The designed DBPs show minimal overlap with nature DBPs, confirming novelty. Most designs exhibit low sequence similarity while retaining good pTM scores. Notably, a cluster with low pTM and low sequence similarity arises due to structure prediction’s reliance on homology, assessing the foldability of such sequences remains challenging.

**Fig. 6.**
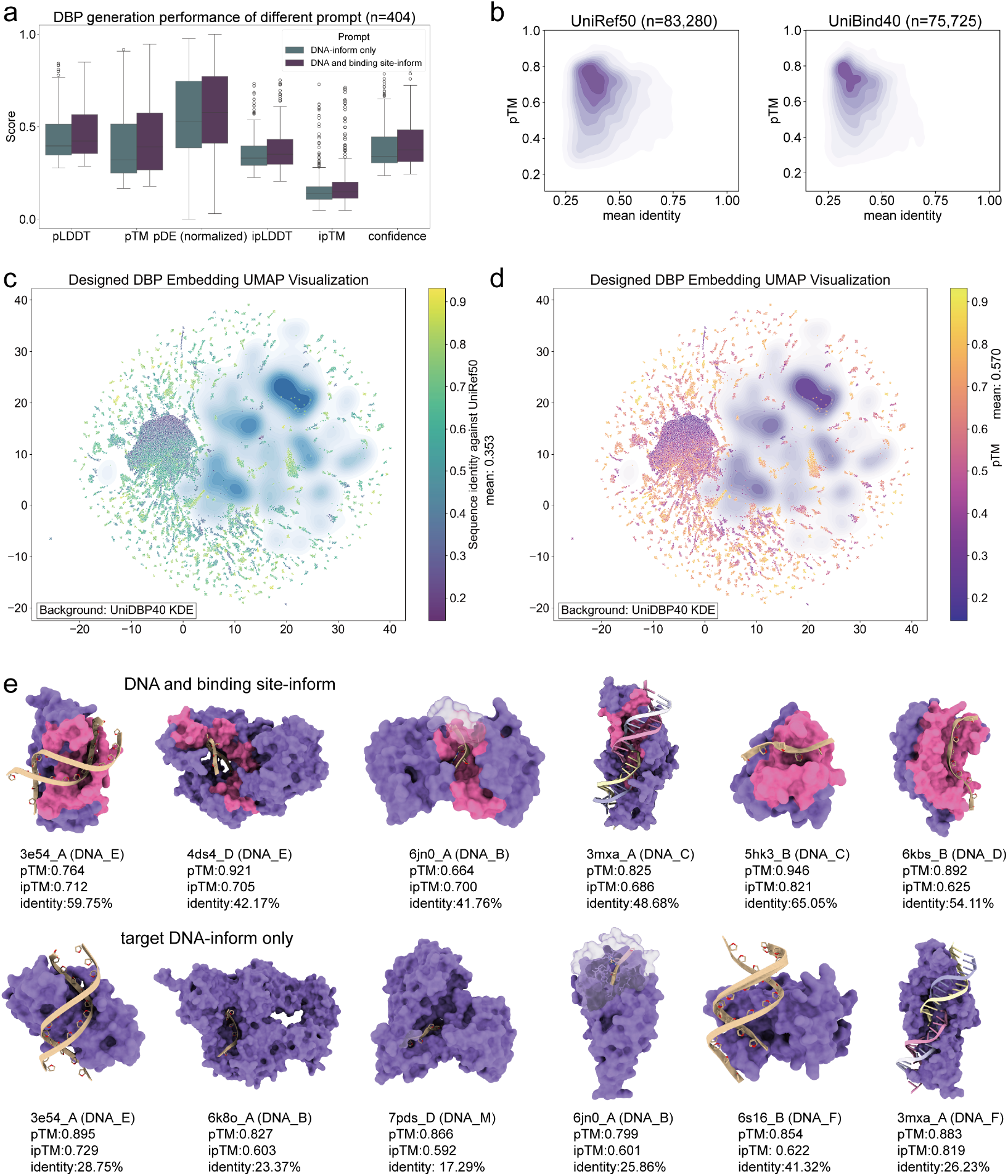
DNA-conditioned binder generation. **a**. Comparison of the generation performance on the DBP test set of different prompts, i.e., DNA and binding site-inform and DNA-inform only. **b**. The kernel density maps between the sequence identity and pTM. For the 101,950 generated sequences, MMseqs2 was used to search the UniRef50 and UniBind40 databases, and similar sequences were found for 83,280 and 75,725 samples, respectively. For each sample, the average sequence identity was taken. **c-d**. UMAP visualization of the embedding of 101,950 generated DBP sequences and UniDBP40 background (170,264 native non-redundant DBPs). **e**. Examples of DBPs generated by different prompts. Prompted residues are highlighted in pink. Complex structures were predicted using boltz. Sequence identity was calculated against the natural sequence of PDB templates.

EiRA fully retains ESM3’s capacity for flexible multimodal input, combining sequence and structural information at any residue. This enables protein generation conditioned solely on DNA by masking protein sequence/structure. Using 404 sample pairs (DBP275 protein sequences and interacting DNA chains from PDB complexes), we performed generation tasks conditioned on: 1) DNA + binding site, and 2) DNA alone. Structural confidence (pTM) decreased only marginally when only DNA was provided, despite the absence of protein information (Fig. 6a). Representative structures for different prompts are shown (Fig. 6e: DNA+binding site above; DNA-only below). Given the binding sites, the target DNA is embedded within its groove (pink highlight). Given only the target DNA, the model generates proteins with pockets accommodating the DNA. Moreover, DNA-only designs exhibit significantly greater sequence divergence from natural binders compared to DNA+binding site designs, providing a powerful method for expanding novel DBP libraries. We performed 100 ns MD simulations on these 6 DNA-given-only generation (Fig. S9 and S10). RMSD and H-bone curves confirmed that not only the designed proteins remain stable, but also maintain dynamic interactions with target DNA. The above findings demonstrate the potential of EiRA to generate binding domains and scaffolds solely from DNA sequences.

### EiRA generates highly divergent mutants

#### 10 mutants of a single Tnpb template

Using Tnpb (Transposase B, a micro RNA-guided programmable DNA endonuclease of length 408 residues; PDB ID: 8exa) as an example, we preserved the amino acid types at 194 key positions (mutation rate: 52.45%, see Table S4), while the remaining residues were masked and generated by EiRA. A total of 500 candidate sequences were input into AF3 along with the reRNA and target DNA, and the top five ranked by pTM were selected for synthesis and expression testing. Previous studies have indicated that removing N consecutive residues within the C-terminal disordered region can enhance Tnpb activity. Accordingly, we deleted 15 positions at the C-terminus and evaluated five truncation mutants following the same procedure as for the full-length variants. Semi-quantitative analysis showed that, among the five full-length mutants, two exhibited higher expression levels than wild-type Tnpb (see Fig. 7b). After C-terminal truncation, four out of the five variants showed expression levels surpassing that of the native template, consistent with earlier reports. Sequence alignment further revealed that these mutants share relatively low similarity with the natural Tnpb family (see Table S4).

**Fig. 7.**
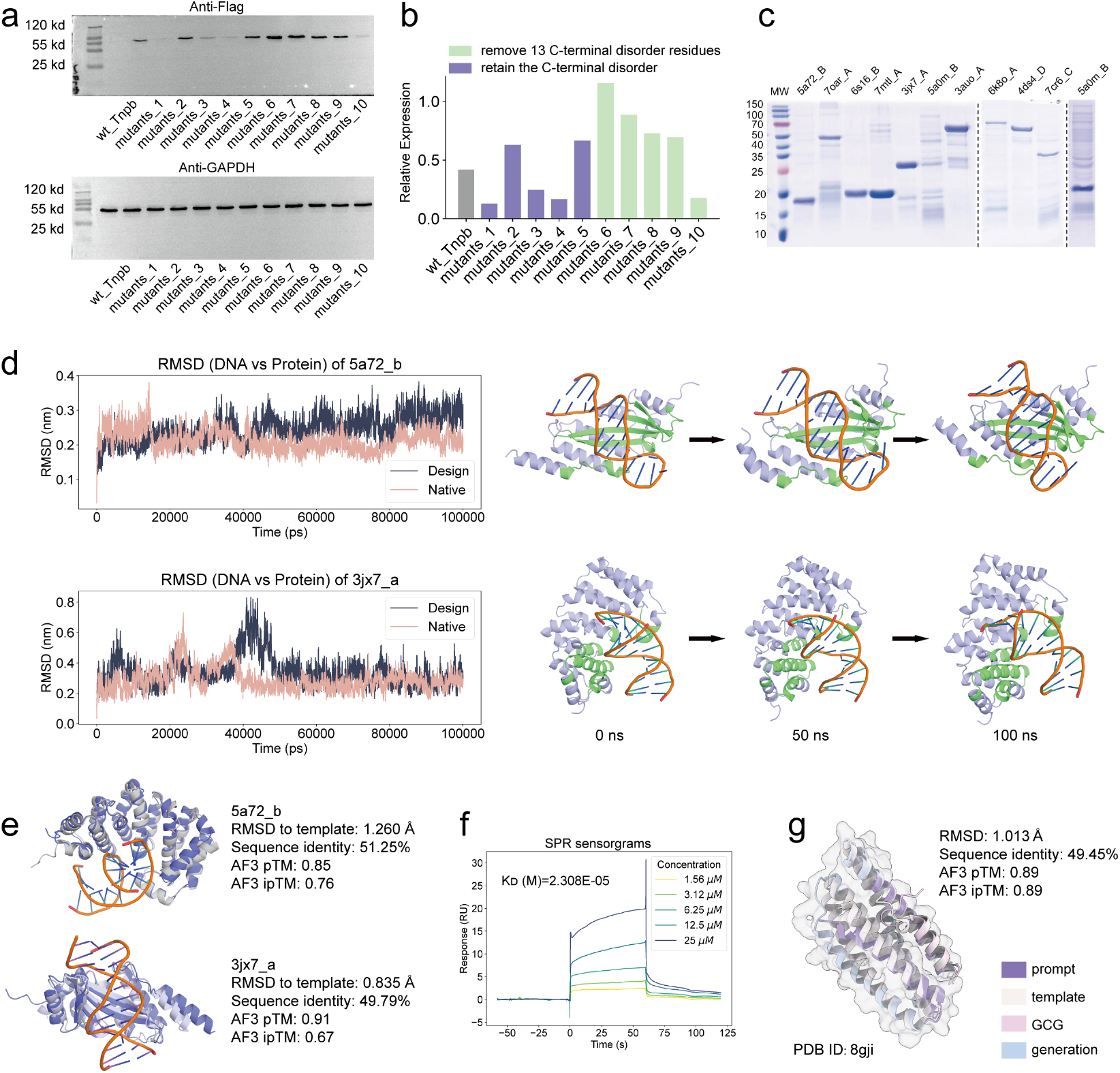
EiRA generates highly divergent protein variants and “one-shot” functional binder design. **(a). WB analysis of TnpB variants**. Expression levels of wild-type (wt), full-length mutants (1–5), and C-terminally truncated mutants (6–10) were detected using Anti-Flag antibodies, with Anti-GAPDH as a loading control; **(b). Quantification of TnpB relative expression using ImageJ** ^**51**^. Green bars indicate C-terminal truncation mutants, which generally exhibit higher expression than full-length variants (purple); **(c). SDS-PAGE analysis of 10 purified DNA-binding protein variants**. The dashed lines indicate splicing of lanes from the same gel; **(d). MD simulation analysis of DBP variants** (5a72_b and 3jx7_a). Left: RMSD curves showing the stability of DNA against the protein for both the EiRA-generated design and the native template over a 100 ns trajectory. Right: Time-lapse structural snapshots (0 ns, 50 ns, and 100 ns) illustrating the conformational evolution and persistent binding interface of the protein-DNA complexes. The prompt residues are highlighted in green; **(e). Structural superposition of designed DBP variants against templates**. Representative cases (5a72_b and 3jx7_a) show high structural fidelity (RMSD < 1.3 Å) to their respective templates despite low sequence identity (∼50%) and robust AF3 confidence scores (pTM/ipTM). **(f). SPR sensorgrams of a functional binder**. Binding interaction between the EiRA-generated binder and the GCG peptide target across a concentration gradient (1.56 to 25 *uM*(, yielding a *K*_*D*_ of 23.08 *uM* ; **(g). Structural superposition of the AF3-predicted EiRA binder (blue) and the GCG peptide (pink) complex against the native template (grey; PDB ID: 8gji)**. The design maintains the native fold (RMSD: 1.013 Å) despite low sequence identity (49.45%). All the mutants were expressed in E. coli and the plasmid structures see Fig. S13. The WB, SPR, and MD experiment details can be found at Supplementary Texts S2, S3, and S4.

These results confirm that EiRA can effectively guide the extensive mutagenesis of natural scaffolds to produce variants with improved expression profiles, even when sequence similarity to the natural family is low.

#### Mutations of 10 DNA-binding proteins

To assess the generalizability of EiRA across the proteome, we extended EiRA to 10 distinct DNA-binding proteins (Table S5). For each template, a single high-scoring EiRA variant was selected for experimental validation. Remarkably, SDS-PAGE analysis confirmed that all 10 designs were successfully expressed and purified (Fig. 7c), despite possessing low sequence identities to their respective templates (Table S3). To further evaluate the structural integrity and binding stability of these variants, we performed 100 ns all-atom MD simulations on these 10 cases. The ubiquitous presence of hydrogen bonds directly demonstrates the interaction (right of Fig. S11). Taking 5a72_b and 3jx7_a (left of Fig. 7d) as examples, the RMSD profiles demonstrated that the DNA-protein complexes reached dynamic equilibrium, with the designed variants exhibiting stability comparable to their native templates. Notably, time-lapse structural snapshots confirmed that the DNA remained securely within the binding pocket throughout the trajectory, maintained by a persistent interfacial network (right of Fig. 7d). High-confidence AF3 predictions further corroborated these results, yielding low RMSD (< 1.3 Å) relative to the templates and robust ipTM scores (Fig. 7e). The successful retrieval of foldable, functional proteins at such high divergence suggests that EiRA has acquired deep structural knowledge, enabling it to navigate the fitness landscape to identify robust sequences far removed from native optima.

### EiRA generates new Glucagon binder

To demonstrate the EiRA’s capacity for generalization beyond monomeric stability to inter-biomolecular interactions, we applied EiRA to design binders for 4 peptide targets (GCG, BCL, PsnA214-38, and Chromatin structure modulator; Table S6). In a single attempt, we successfully expressed and purified a “one-shot” Glucagon peptide binder (Template PDB ID: 8gji), achieving a purity of 95.9% (Fig. S12).

Structural assessment using AF3 predicted a high-confidence complex (pTM and ipTM scores of 0.89; Fig. 7g). The generated binder adopts a well-defined helical conformation that superimposes closely with the reference template (RMSD = 1.013 Å). Crucially, this structural conservation is achieved despite a sequence identity of only 49.45%. This substantial divergence highlights EiRA’s capability to generalize beyond the training distribution; rather than merely recapitulating native sequences, the model explores novel regions of sequence space to identify solutions that satisfy the precise geometric constraints required for binding. We experimentally characterized the binding affinity using SPR. The EiRA-designed binder exhibited clear, dose-dependent binding kinetics to the GCG target with a dissociation constant *K*_*D*_ of 23.08 *uM* (Fig. 7f). This validated binder serves as a promising lead scaffold for the development of therapeutic glucagon antagonists targeting hyperglycemia in diabetes and metabolic disorders.

Overall, the convergence of high-confidence structural prediction and confirmed biophysical interaction underscores EiRA’s potential in generating functional binders with specific affinity for peptide targets. Novel binder with low sequence identity demonstrates that EiRA decouples structural integrity from sequence homology, enabling the generation of hyper-mutated yet functional protein variants.

## Discussion

In this study, we introduce EiRA, a multimodal protein language model specifically designed for biomolecular-binding protein generation. Built upon the general foundation model ESM3, EiRA undergoes two rounds of post-training: domain-adaptive masking training and binding site-informed preference optimization. The main contributions of this work are summarized as follows:

1. We curated UniBind40, a large-scale dataset of biomolecular-binding proteins, to support customized self-supervised training;
2. Comprehensive multi-dimensional evaluations demonstrate that EiRA outperforms the native ESM3s (1.4 billion) and ESM3m (7 billion), and is comparable with ESM3l (98 billion), across multiple ligand-binding protein design tasks;
3. The optimized loss function effectively alleviates ESM3’s propensity for repetitive generation, leading to improved design quality;
4. EiRA’s representations facilitate direct predictions for diverse biomolecular binding-related downstream tasks and offer superior protein characterization compared to the native ESM3s;
5. By integrating DNA sequence, EiRA supports the design of DNA-binding proteins under DNA conditions, thereby broadening the design paradigm;
6. We achieved the “one-shot” design of a highly divergent GCG peptide binder with micromolar affinity.

We summarized 4 challenges for AI protein design, hoping to provide potential insights for future studies.

1. Constructing a high-quality protein-complex distillation dataset to compensate scarce natural complex;
2. Developing screening strategies integrated with biological context and microenvironment to bridge the gap between computational design and practical application;
3. Establishing alternative screening methods for long and orphan proteins, where conventional structure prediction remains unreliable;
4. Balancing the inherent trade-off between diversity and reliability of the generated protein.

Compared with the closed-source ESM3m and ESM3l, we fully opened up the dataset, model weights, training, and inference script for freely academic use. Although there is still room for improvement, the proposed EiRA has the potential to become a powerful protein design tool, thereby accelerating the development of biomedicine, such as gene editing, immunotherapy and drug discovery.

## Data Availability

The data, source code, and standalone program of EiRA are freely accessible at https://github.com/pengsl-lab/EiRA. The pretraining data set and the weight of all EiRA models are available for academic use at https://huggingface.co/zengwenwu/EiRA/tree/main.

## Author Contributions

Wenwu Zeng: Conceptualization, Methodology, Software, Visualization, Writing original draft. Haitao Zou: Methodology. Xiaoyu Li: Visualization. Yutao Dou: Software. Xiaoqi Wang: Conceptualization, Writing-review & editing. Shaoliang Peng: Funding acquisition, Resources, Supervision.

## Acknowledgment

This work was supported by NSFC-FDCT Grants 62361166662; NSFC Grants 625B2068; National Key R&D Program of China 2023YFC3503400, 2022YFC3400400; The Innovative Research Group Project of Hunan Province 2024JJ1002; Hunan Science and Technology Innovation Plan 2025ZYJ003; Key R&D Program of Hunan Province 2023GK2004, 2023SK2059, 2023SK2060; Top 10 Technical Key Project in Hunan Province 2023GK1010; Key Technologies R&D Program of Guangdong Province (2023B1111030004 to FFH); Postgraduate Research Innovation Project of Hunan Province CX20250637. We would like to thank the Fund of the National Supercomputing Center in Changsha (http://nscc.hnu.edu.cn/), Peng Cheng Lab, Key Laboratory of High-Performance Distributed Ledger Technology and Digital Finance (Ministry of Education), and Hunan Research Center of the Basic Discipline for Cell Signaling.

Thanks to EvolutionaryScale for developing the ESM3 foundational language model and ESMFold. Thanks to DeepMind for developing the AlphaFold3 and the MIT Jameel Clinic for reproducing the Boltz.

## Supplementary Text S1: Feature generation

For each protein, the ESM3 feature representation is applied. The DSSP is utilized to calculate eight protein secondary structure (SS) states. The solvent accessibility (SA) is calculated by the Shrake-Rupley algorithm in the Biotite package. We downloaded the InterPro annotations of all UniProtKB sequences at https://ftp.ebi.ac.uk/pub/databases/interpro/current_release/protein2ipr.dat.gz and then extracted the corresponding UniBind40 records from them. For BioDPO and all test sets, only sequences and atomic coordinates are utilized as prompts. All these feature sources are fed into the ESM3 tokenizer to form numerical tokens for training.

## Supplementary Text S2: Western Blot

Recombinant proteins expressed in E. coli were extracted using lysis buffer supplemented with PMSF and a protease inhibitor cocktail. Protein concentration was determined using a BCA kit (Beyotime). Samples (100 *u*g) were separated by SDS-PAGE and transferred onto PVDF membranes (Millipore). After blocking with 5% non-fat milk, membranes were incubated overnight at 4°C with anti-Flag (1:5,000, Proteintech) or anti-GAPDH (1:50,000, Proteintech) antibodies. Blots were subsequently incubated with HRP-conjugated Goat Anti-Mouse IgG secondary antibody (1:10,000, Proteintech) and visualized using an ECL substrate system.

## Supplementary Text S3: Surface Plasmon Resonance

Surface Plasmon Resonance (SPR) analysis was performed on a CM5 sensor chip, where the ligand protein was immobilized to approximately 2200 response units (RU) using a concentration of 5 *u*g/mL in sodium acetate buffer (pH 4.0). Serial dilutions of the analyte peptide (0 to 25 *u*M) were injected over the surface with key parameters set as follows: a contact time of 60 seconds, a constant flow rate of 30 *u*L/min, a dissociation time of 60 seconds, and a temperature of 25°C, using HBS-EP+ as the running buffer.

## Supplementary Text S4: Molecular dynamics simulation

We performed all-atom molecular dynamics simulations using the GROMACS tool with the “amber14sb_parmbsc1” force field, which incorporates refined parameters for protein side chains and DNA *α*/*γ* backbone torsions. The complex was solvated in a dodecahedron periodic box with TIP3P water molecules, ensuring a minimum distance of 1.0 nm between the solute and the box edges, and neutralized with 0.15 M NaCl to mimic physiological ionic strength.The simulation protocol initiated with a 50,000-step energy minimization using the steepest descent algorithm (emtol = 1000.0 *kJ* · *mol*^−1^ · *nm*^−1^), followed by a 100 ps NVT equilibration and a 100 ps NPT equilibration, both employing the V-rescale thermostat at 300 K. Pressure was maintained at 1.0 bar during NPT and production phases using the Parrinello-Rahman barostat with an isothermal compressibility of 4.5 × 10^−5^*bar*^−1^. Long-range electrostatic interactions were treated via the Particle Mesh Ewald method with a 1.0 nm cutoff, while van der Waals forces were managed using a Verlet cutoff scheme at 1.0 nm. All bonds involving hydrogen atoms were constrained using the LINCS algorithm, allowing for an integration time step of 2 fs. A 100 ns production run was executed, with trajectories recorded every 10 ps. Post-simulation analysis involved PBC correction and centering, followed by the calculation of RMSD and interfacial hydrogen bonding frequency to characterize the binding interface.

**Fig. S1.**
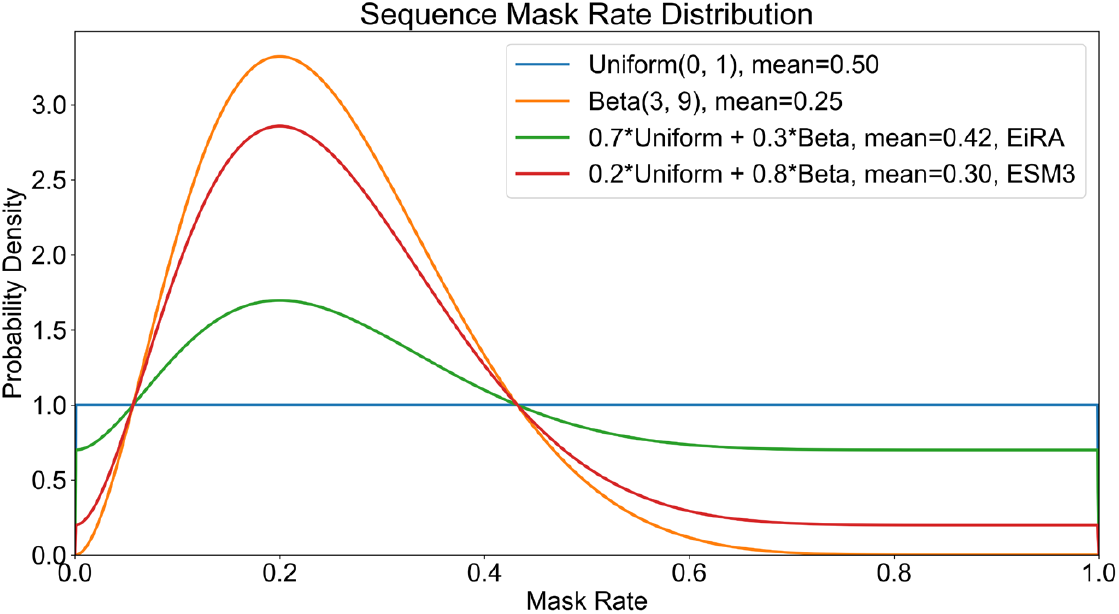
EiRA adopts a higher sequence mask rate than that of ESM3.

**Fig. S2.**
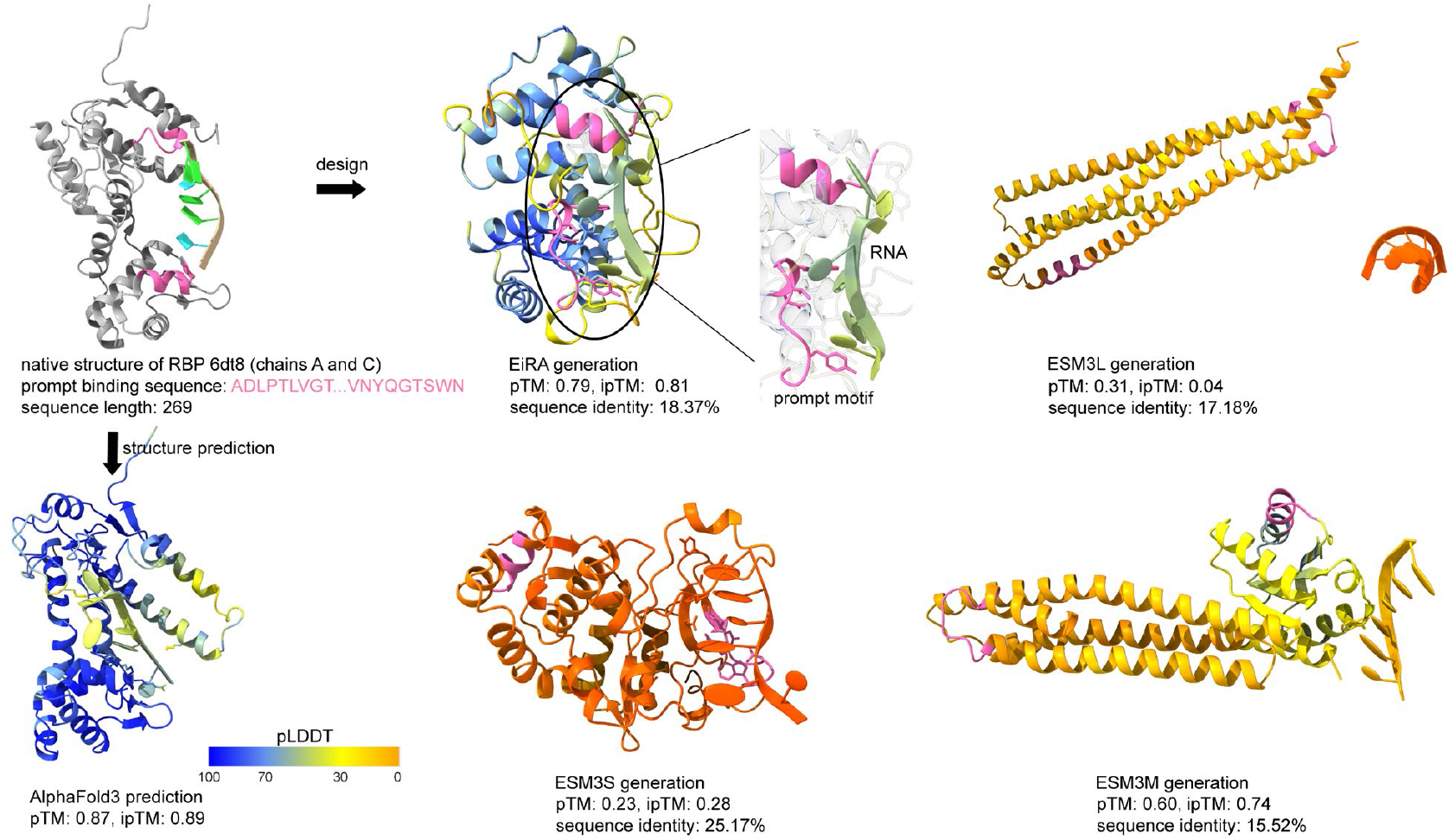
Case study on a RBP (PDB ID: 6dt8, Bacteriophage N4 RNA polymerase II elongation complex 1). The new protein (a sequence identity of 18.37% against PDB template) designed by EiRA maintains high stability while retaining the ability of the template motif to contact RNA ligands. In contrast, all three versions of ESM3 produced low-confidence proteins. The protein sequences were designed by four MPLMs from the template motif (highlight with pink) and then the complex structures were predicted by Boltz.

**Fig. S3.**
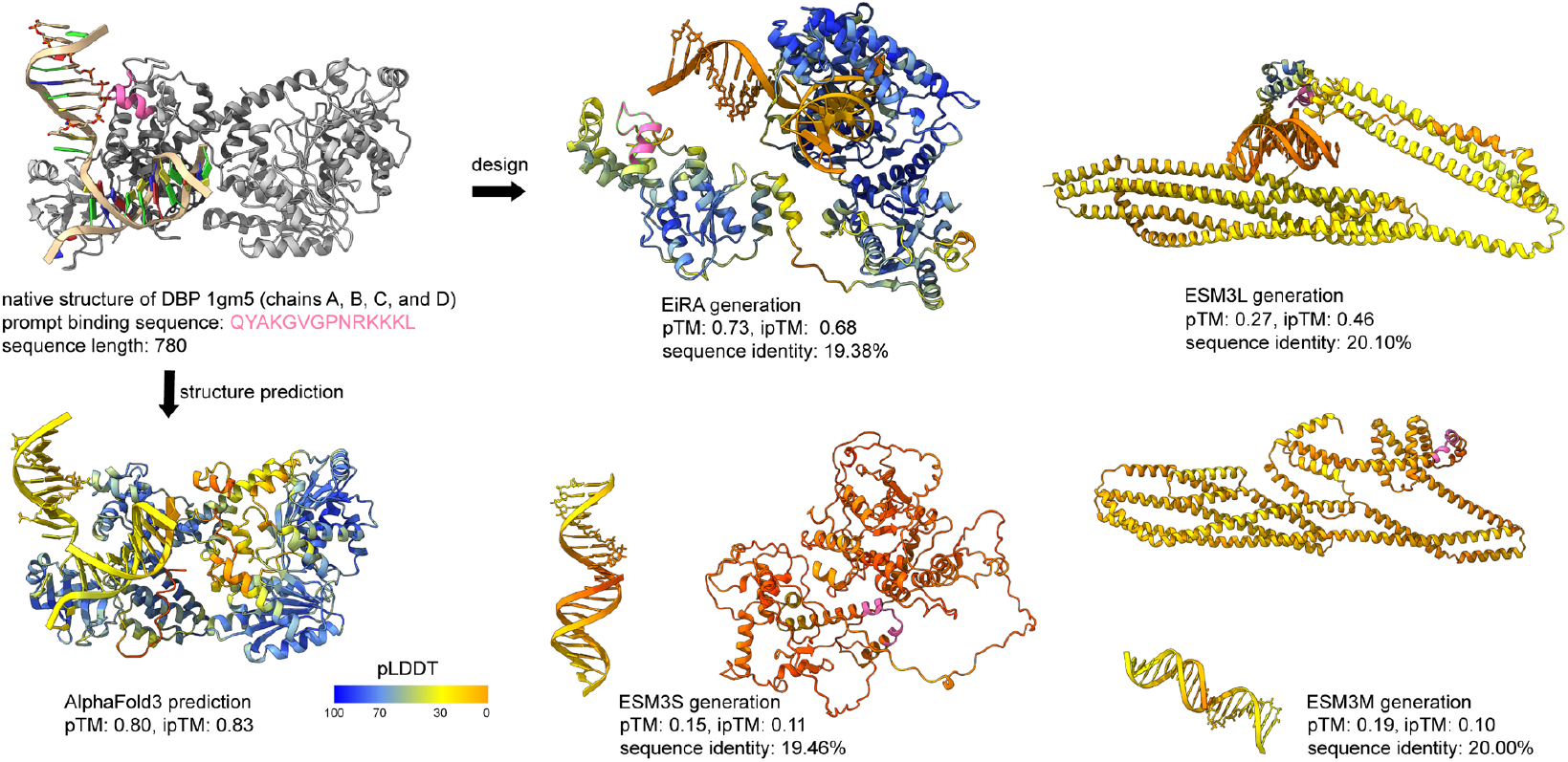
Case study on a DBP (PDB ID: 1gm5, RecG protein, DNA-specific helicase). The novel proteins generated by EiRA have high confidence and are predicted to interact with DNA, while the confidence of the other three is low, and the ones generated by ESM3s and ESM3m are predicted not to bind DNA.

**Fig. S4.**
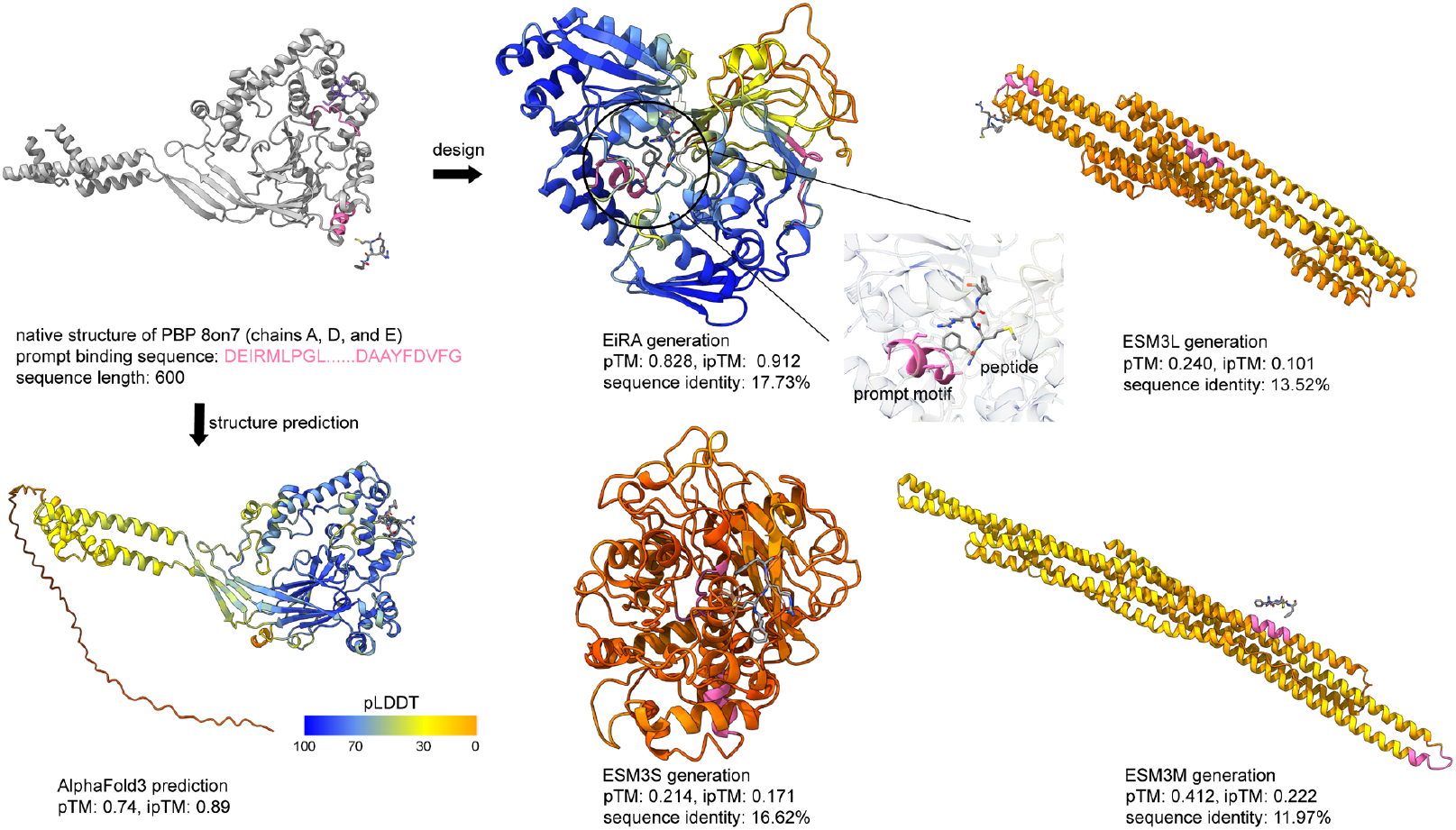
Case study on a PBP (PDB ID: 8on7, Sort-Tagged Drosophila Cryptochrome). The pTM and ipTM scores of EiRA designed protein are higher than that of the AF3 structure of native sequence.

**Fig. S5.**
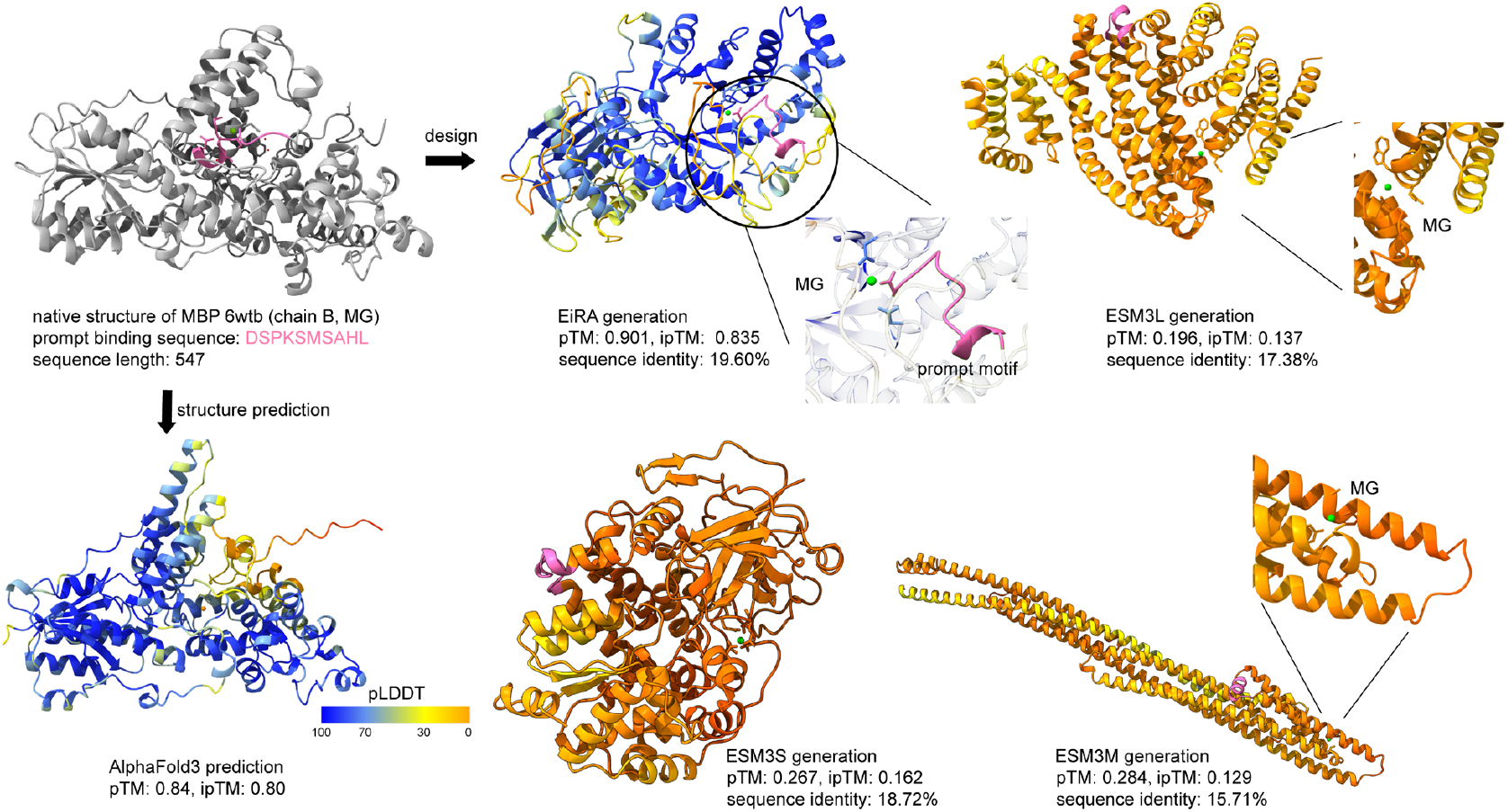
Case study on a MBP (PDB ID: 6wtb, FMRFa-bound Malacoceros FaNaC1).

**Fig. S6.**
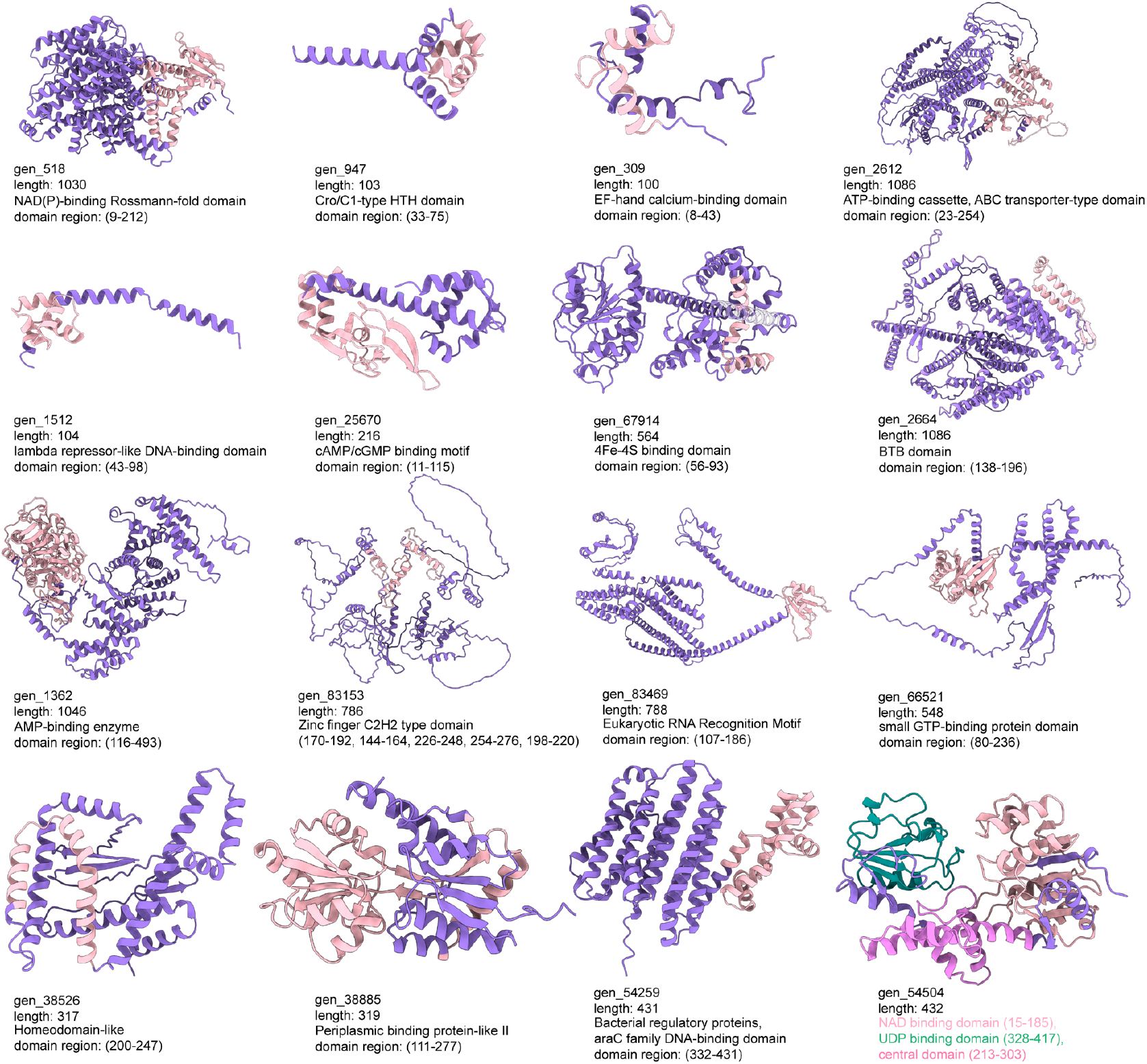
EiRA generates various biomolecule binding domains unconditionally. The samples are selected from 100,000 unconditionally generated sequences, and AF3 is used to predict the tertiary structure. The domain annotations are predicted using InterProScan tool.

**Fig. S7.**
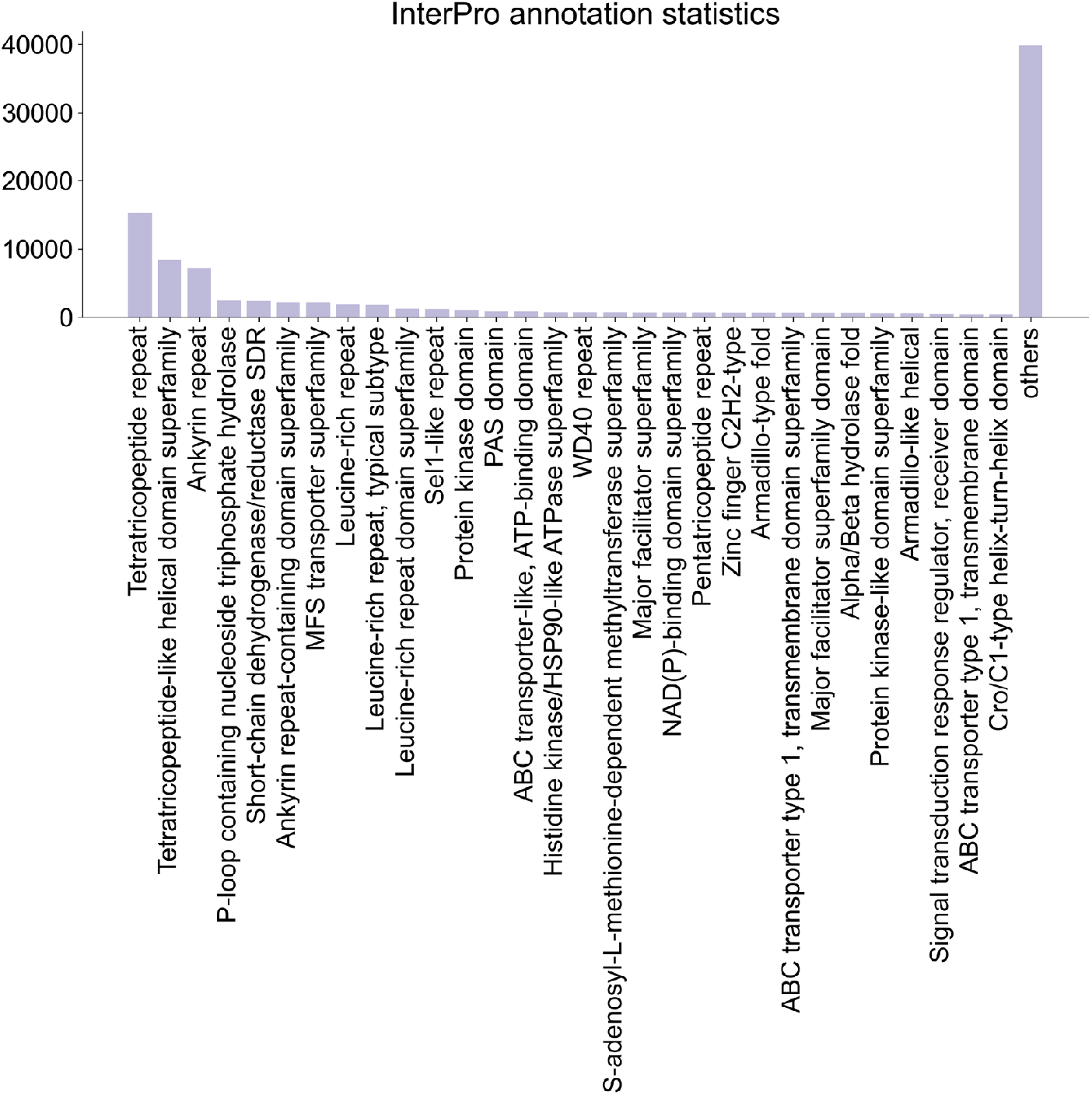
InterProScan records distribution of 100,000 unconditional generated sequences (no redundancy was removed).

**Fig. S8.**
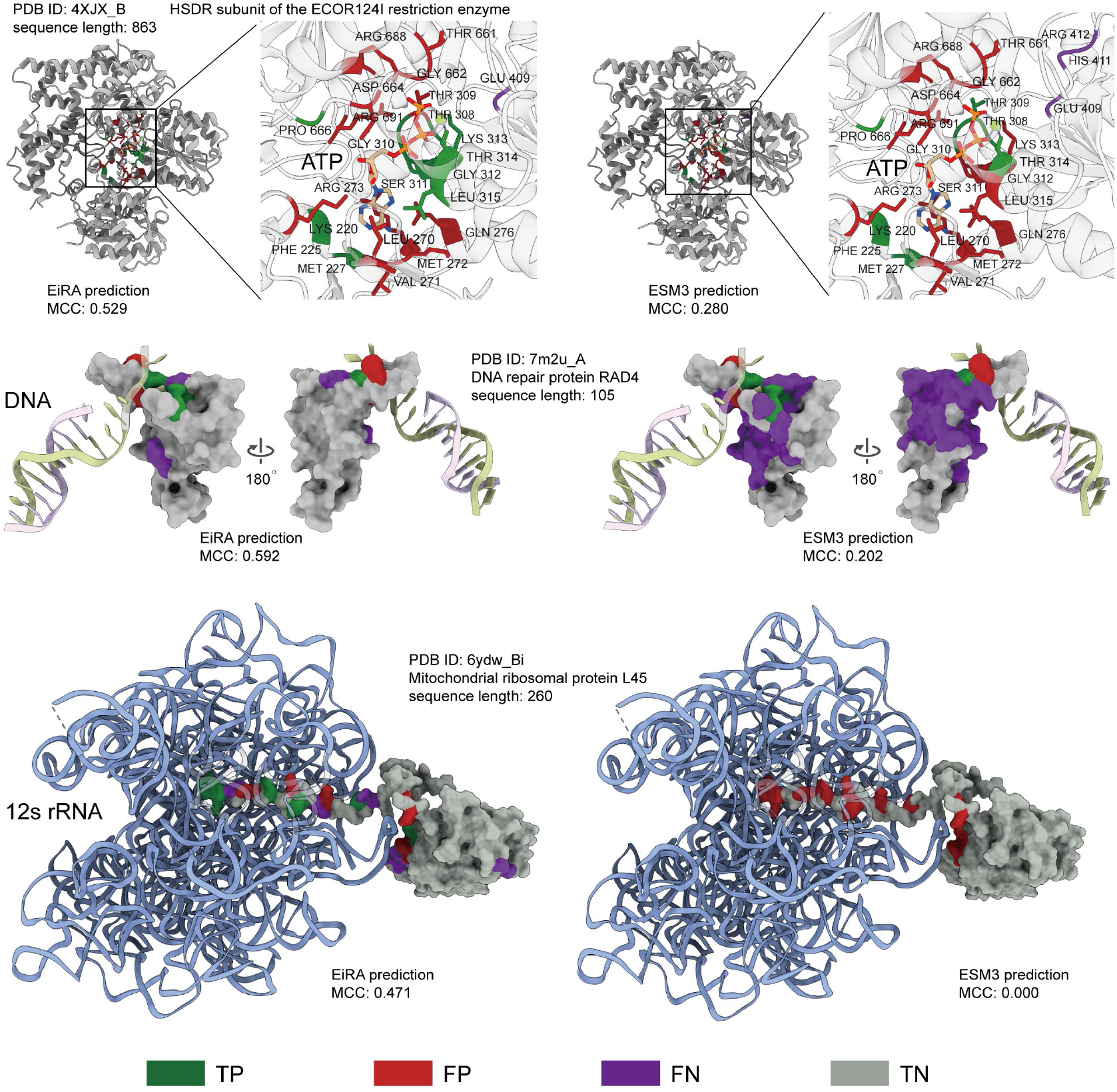
Case studies of DNA-, RNA-, and ATP-binding interface predictions.

**Fig. S9.**
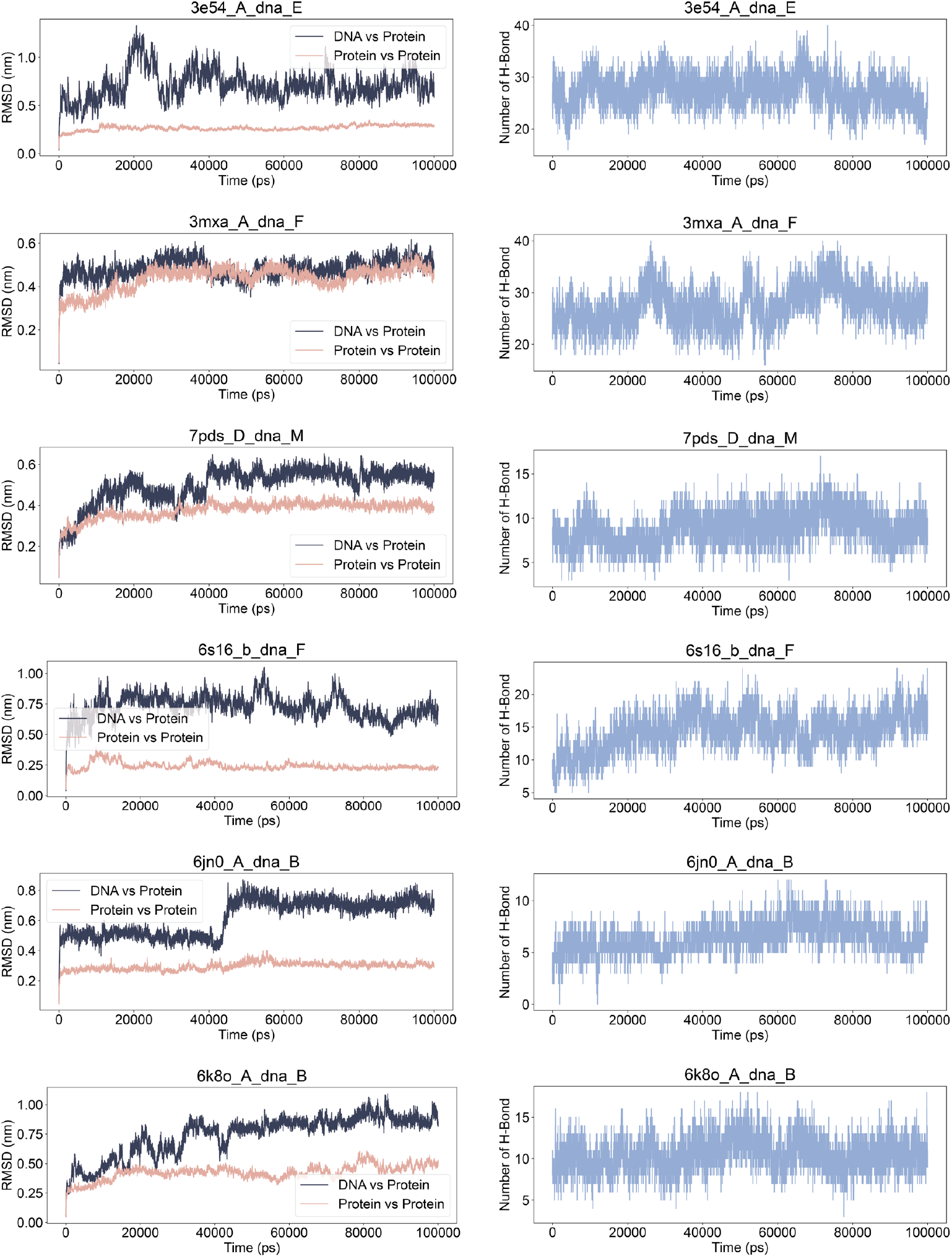
RMSD and H-Bond of MD simulations on 6 DBPs generated by EiRA with DNA sequence prompt only.

**Fig. S10.**
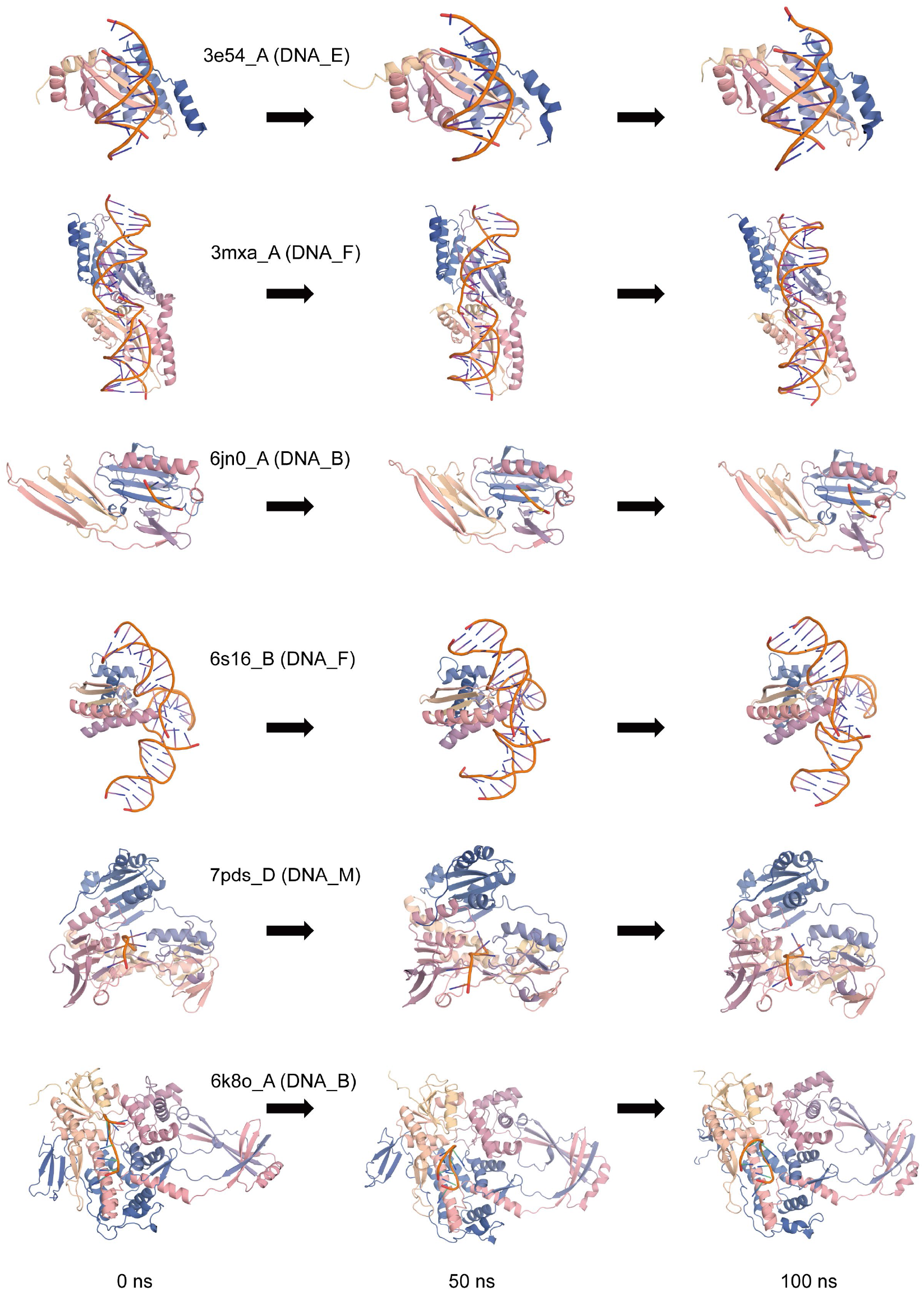
Visualization of MD simulations on 6 DBPs generated by EiRA with DNA sequence prompt only.

**Fig. S11.**
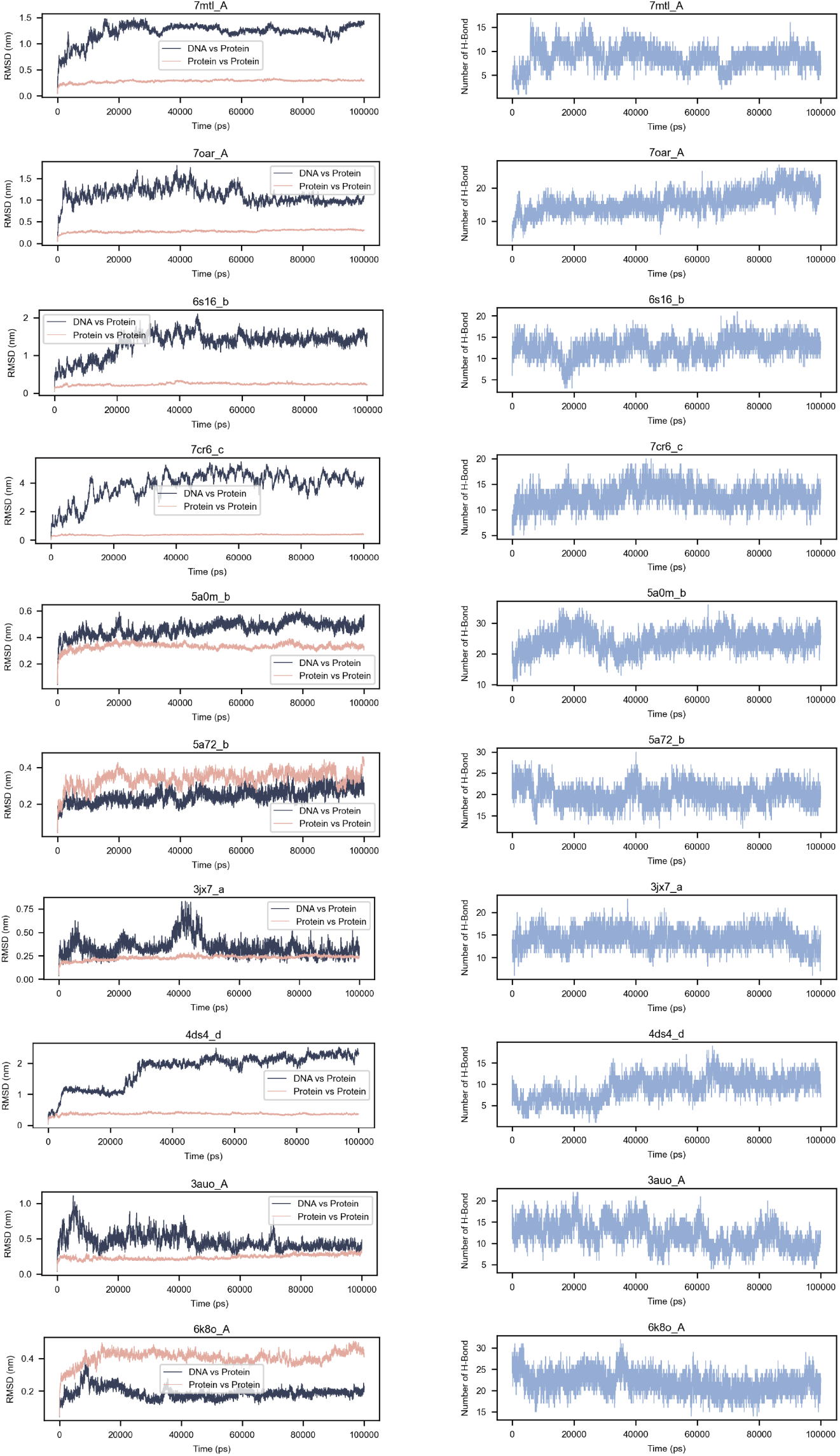
RMSD and H-Bond of MD simulations on 10 purified DBPs.

**Fig. S12.**
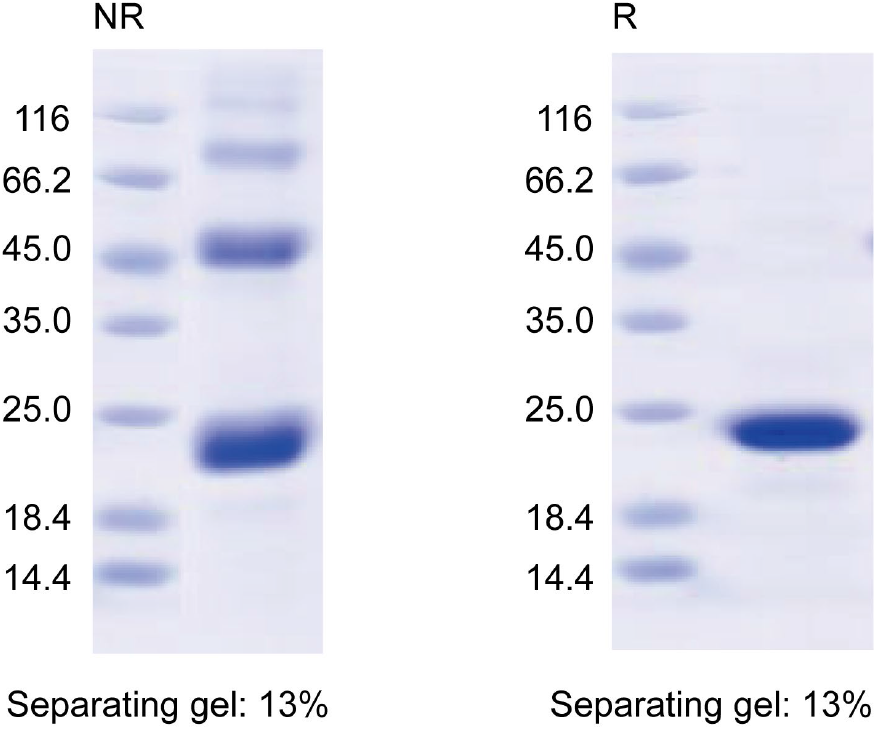
SDS-PAGE detection of the designed GCG binder (Purity: 95.9%).

**Fig. S13.**
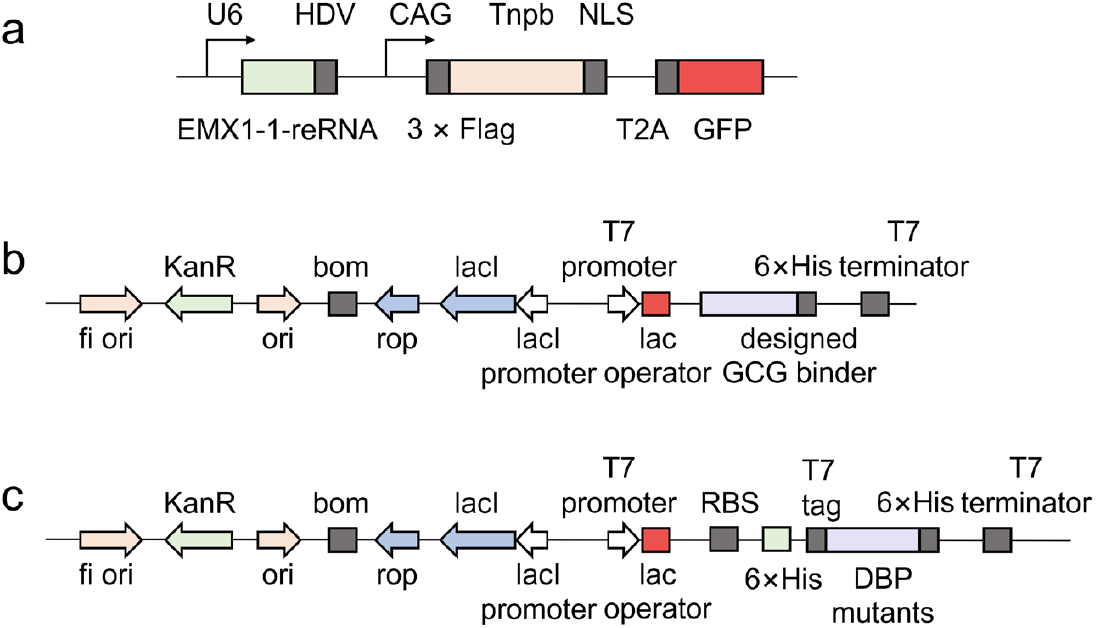
The plasmid structure of the recombinant protein. **a**. Tnpb mutants for determination of expression; **b**. GCG binder for SPR. **c**. Mutants of DNA-binding protein.

**Table S1.** The impact of updatable Transformer blocks on performance.

| Structure predictor | Evaluation index | Fine-tuning layers | DBP | RBP | PBP | ReBP | MBP | PPI | MDBP | MRBP |
| --- | --- | --- | --- | --- | --- | --- | --- | --- | --- | --- |
| ESM3 | pTM | 0 (ESM3) | 0.503 | 0.502 | 0.540 | 0.630 | 0.530 | 0.439 | **0.442** | 0.516 |
|  |  | 16 | 0.502 | 0.496 | 0.559 | 0.656 | 0.578 | 0.445 | 0.406 | 0.498 |
|  |  | 24 | 0.453 | 0.487 | 0.519 | 0.600 | 0.511 | 0.415 | 0.383 | 0.472 |
|  |  | 32 | **0.506** | **0.534** | 0.563 | **0.657** | **0.580** | **0.462** | 0.431 | **0.532** |
|  |  | 48 | 0.485 | 0.529 | **0.566** | 0.654 | 0.572 | 0.461 | 0.430 | 0.517 |
|  | pLDDT | 0 (ESM3) | 0.700 | 0.691 | 0.708 | 0.742 | 0.680 | 0.681 | 0.679 | 0.727 |
|  |  | 16 | 0.742 | 0.722 | **0.780** | **0.802** | **0.775** | 0.732 | 0.722 | 0.746 |
|  |  | 24 | 0.680 | 0.683 | 0.719 | 0.732 | 0.694 | 0.719 | 0.709 | 0.726 |
|  |  | 32 | **0.745** | 0.727 | 0.764 | 0.787 | 0.751 | 0.733 | 0.729 | 0.752 |
|  |  | 48 | 0.744 | **0.738** | 0.772 | 0.794 | **0.775** | **0.769** | **0.759** | **0.775** |
| ESMFold | pTM | 0 (ESM3) | 0.418 | 0.429 | 0.396 | 0.548 | 0.444 | 0.363 | 0.374 | 0.317 |
|  |  | 16 | **0.443** | **0.434** | **0.449** | **0.603** | **0.528** | 0.397 | **0.405** | 0.473 |
|  |  | 24 | 0.387 | 0.405 | 0.377 | 0.509 | 0.410 | 0.345 | 0.343 | 0.430 |
|  |  | 32 | 0.416 | 0.426 | 0.419 | 0.570 | 0.477 | **0.398** | 0.396 | **0.480** |
|  |  | 48 | 0.380 | 0.392 | 0.425 | 0.564 | 0.479 | 0.389 | 0.388 | 0.461 |
|  | pLDDT | 0 (ESM3) | 0.441 | 0.437 | 0.452 | 0.535 | 0.453 | 0.469 | 0.524 | 0.448 |
|  |  | 16 | **0.504** | 0.492 | **0.521** | **0.601** | **0.557** | 0.563 | 0.616 | 0.626 |
|  |  | 24 | 0.440 | 0.449 | 0.443 | 0.512 | 0.443 | 0.462 | 0.485 | 0.552 |
|  |  | 32 | 0.502 | **0.494** | 0.496 | 0.584 | 0.524 | **0.599** | **0.630** | **0.655** |
|  |  | 48 | 0.485 | 0.484 | 0.512 | 0.583 | 0.531 | 0.584 | 0.605 | 0.617 |
|  | pAE | 0 (ESM3) | 19.83 | 19.54 | 19.121 | 15.92 | 18.94 | 23.80 | 23.47 | 26.47 |
|  |  | 16 | **19.36** | 19.57 | **17.98** | **14.14** | **16.45** | **22.87** | 22.83 | 21.71 |
|  |  | 24 | 20.80 | 20.31 | 19.88 | 17.03 | 19.79 | 24.38 | 24.91 | 23.13 |
|  |  | 32 | 19.97 | **19.50** | 18.61 | 15.01 | 17.72 | 22.94 | **22.81** | **21.36** |
|  |  | 48 | 20.60 | 20.17 | 18.46 | 15.13 | 17.77 | 22.94 | 22.97 | 21.75 |

**Table S2.** The impact of fine-tuning strategy on performance.

| Structure predictor | Evaluation index | Fine-tuning strategy | DBP | RBP | PBP | ReBP | MBP | PPI | MDBP | MRBP |
| --- | --- | --- | --- | --- | --- | --- | --- | --- | --- | --- |
| ESM3 | pTM | base <sup>a</sup> | 0.463 | 0.484 | 0.536 | 0.602 | 0.529 | 0.432 | 0.370 | 0.415 |
|  |  | vanilla LoRA <sup>b</sup> | **0.502** | 0.496 | **0.559** | **0.656** | **0.578** | 0.445 | 0.406 | 0.498 |
|  |  | LoRA+ <sup>c</sup> | 0.471 | **0.501** | 0.557 | 0.641 | 0.569 | 0.446 | 0.411 | **0.512** |
|  |  | LoKr <sup>d</sup> | 0.421 | 0.459 | 0.504 | 0.596 | 0.524 | 0.394 | 0.365 | 0.427 |
|  |  | hydraLoRA <sup>e</sup> | 0.475 | 0.493 | 0.551 | 0.623 | 0.549 | **0.453** | **0.427** | 0.504 |
|  |  | AdaLoRA <sup>f</sup> | 0.449 | 0.477 | 0.518 | 0.612 | 0.523 | 0.406 | 0.373 | 0.430 |
|  | pLDDT | base | 0.667 | 0.663 | 0.684 | 0.707 | 0.667 | 0.707 | 0.678 | 0.691 |
|  |  | vanilla LoRA | **0.742** | **0.722** | **0.780** | **0.802** | **0.775** | 0.732 | 0.722 | 0.746 |
|  |  | LoRA+ | 0.723 | 0.715 | 0.755 | 0.779 | 0.752 | 0.741 | **0.733** | **0.754** |
|  |  | LoKr | 0.681 | 0.681 | 0.730 | 0.755 | 0.725 | 0.668 | 0.652 | 0.683 |
|  |  | hydraLoRA | 0.688 | 0.677 | 0.754 | 0.767 | 0.748 | **0.752** | 0.673 | 0.715 |
|  |  | AdaLoRA | 0.688 | 0.682 | 0.725 | 0.755 | 0.700 | 0.671 | 0.636 | 0.656 |
| ESMFold | pTM | base | 0.359 | 0.354 | 0.349 | 0.468 | 0.381 | 0.336 | 0.305 | 0.344 |
|  |  | vanilla LoRA | **0.443** | **0.434** | **0.449** | **0.603** | **0.528** | **0.397** | **0.405** | 0.473 |
|  |  | LoRA+ | 0.418 | 0.417 | 0.419 | 0.565 | 0.491 | 0.385 | 0.400 | **0.475** |
|  |  | LoKr | 0.394 | 0.408 | 0.388 | 0.540 | 0.467 | 0.357 | 0.363 | 0.428 |
|  |  | hydraLoRA | 0.381 | 0.389 | 0.400 | 0.552 | 0.451 | 0.369 | 0.359 | 0.459 |
|  |  | AdaLoRA | 0.416 | 0.428 | 0.393 | 0.543 | 0.437 | 0.356 | 0.339 | 0.399 |
|  | pLDDT | base | 0.426 | 0.413 | 0.432 | 0.480 | 0.423 | 0.487 | 0.494 | 0.495 |
|  |  | vanilla LoRA | **0.504** | **0.492** | **0.521** | **0.601** | **0.557** | **0.563** | **0.616** | **0.626** |
|  |  | LoRA+ | 0.484 | 0.475 | 0.496 | 0.567 | 0.524 | 0.537 | 0.599 | 0.619 |
|  |  | LoKr | 0.444 | 0.445 | 0.459 | 0.548 | 0.499 | 0.516 | 0.558 | 0.591 |
|  |  | hydraLoRA | 0.424 | 0.420 | 0.474 | 0.550 | 0.488 | 0.519 | 0.510 | 0.586 |
|  |  | AdaLoRA | 0.456 | 0.460 | 0.471 | 0.550 | 0.470 | 0.500 | 0.507 | 0.540 |
|  | pAE | base | 21.27 | 21.42 | 20.55 | 18.21 | 20.52 | 24.52 | 25.50 | 25.51 |
|  |  | vanilla LoRA | **19.36** | **19.57** | **17.98** | **14.14** | **16.45** | **22.87** | **22.83** | 21.71 |
|  |  | LoRA+ | 19.96 | 20.03 | 18.79 | 15.29 | 17.56 | 23.22 | 22.89 | **21.63** |
|  |  | LoKr | 20.66 | 20.34 | 19.58 | 16.13 | 18.22 | 24.44 | 24.35 | 23.28 |
|  |  | hydraLoRA | 20.64 | 20.48 | 19.14 | 15.74 | 18.56 | 23.52 | 24.21 | 22.03 |
|  |  | AdaLoRA | 19.98 | 19.67 | 19.40 | 16.02 | 19.04 | 24.36 | 25.02 | 24.21 |

- a.* Update the original weights directly, rather than applying adapters and introducing additional parameters. *b.* Low-rank matrix adaptation constructed by multiplying two miniature matrices. *c.* LoRA+ sets different learning rates for the matrices A and B of LoRA. *d.* LoKr employs Kronecker product decomposition. *e.* Asymmetric hybrid expert LoRA, containing a shared matrix A, multiple matrices B, and a router. *f.* AdaLoRA dynamically adjusts the rank of low-rank matrices based on parameter importance scores, optimizing parameter allocation.

**Table S3.** The Summary statistics for the 10 DNA-binding variants, showing the template PDB ID, mutation rate relative to the wild-type, and final purified concentration.

| Template ID | Mutation rate (%) | Concentration (mg/ml) |
| --- | --- | --- |
| 6k8o_A | 64.21 | 0.30 |
| 5a72_B | 48.73 | 0.12 |
| 4ds4_D | 58.29 | 0.39 |
| 7oar_A | 57.69 | 0.10 |
| 6s16_B | 41.82 | 0.15 |
| 7mtl_A | 66.67 | 0.20 |
| 3jx7_A | 51.50 | 0.12 |
| 5a0m_b | 73.68 | 1.05 |
| 7cr6_C | 77.30 | 0.50 |
| 3auo_A | 52.36 | 0.15 |

**Table S4.**
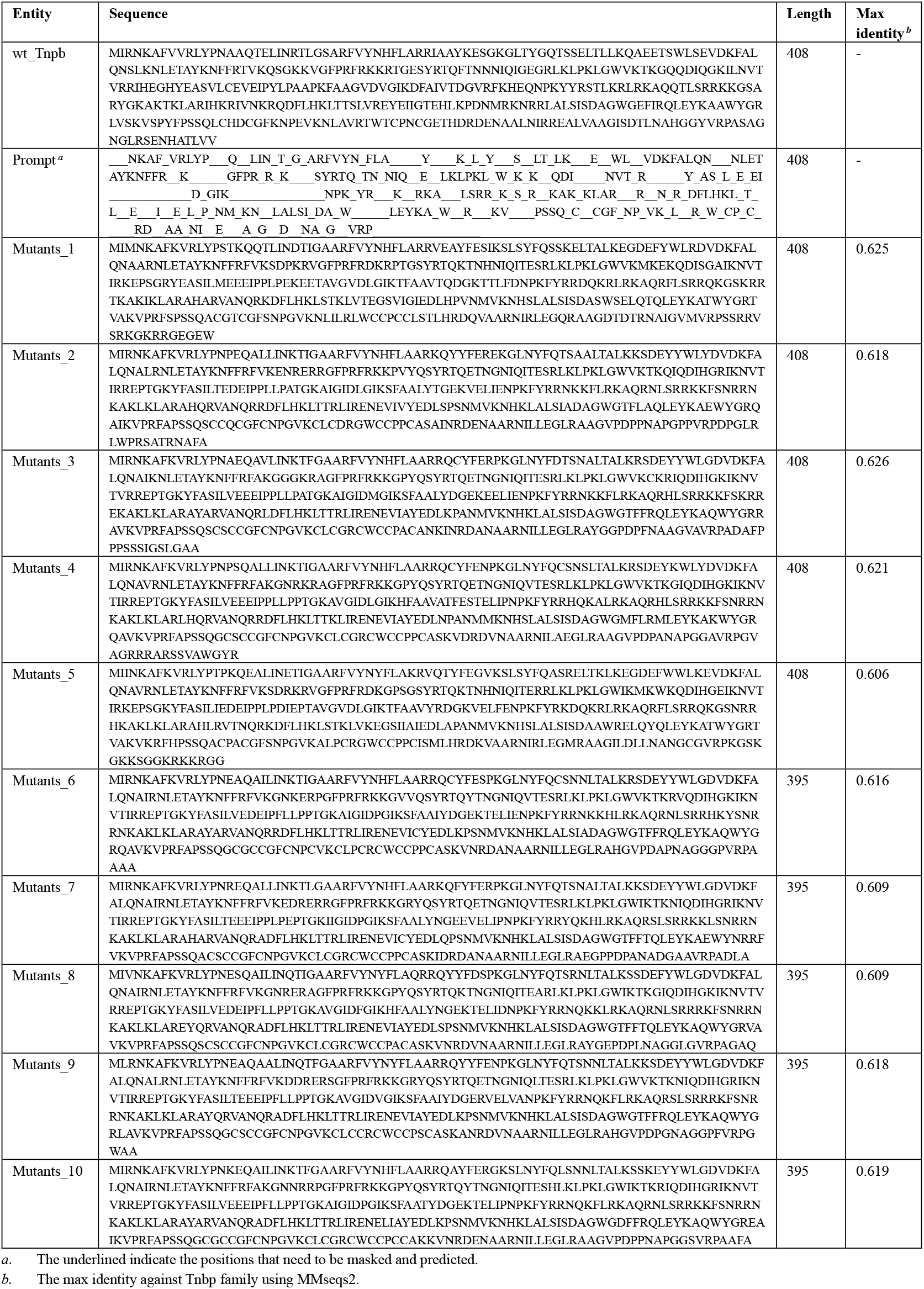
Tnpb sequences for expression detection.

**Table S5.**
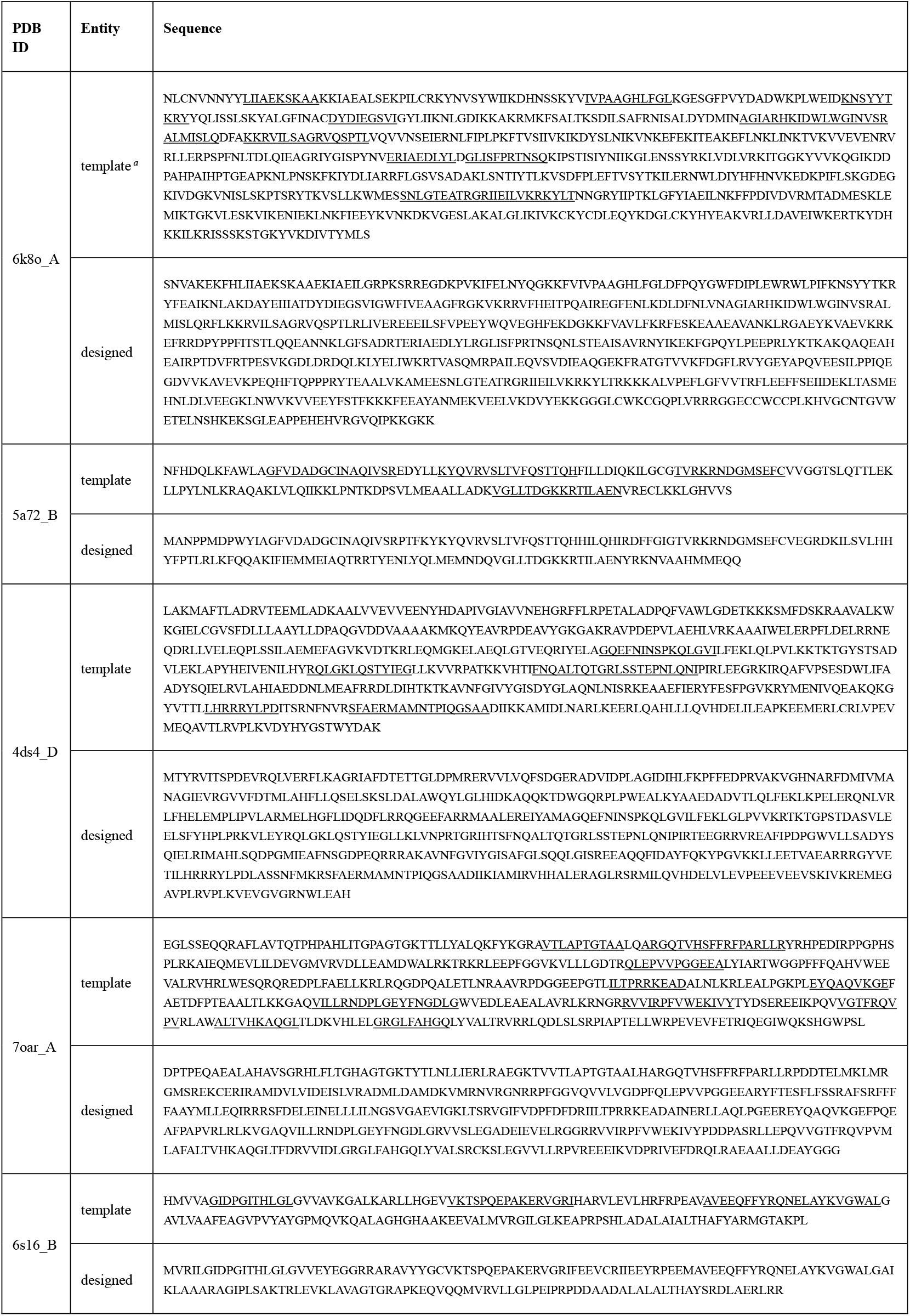
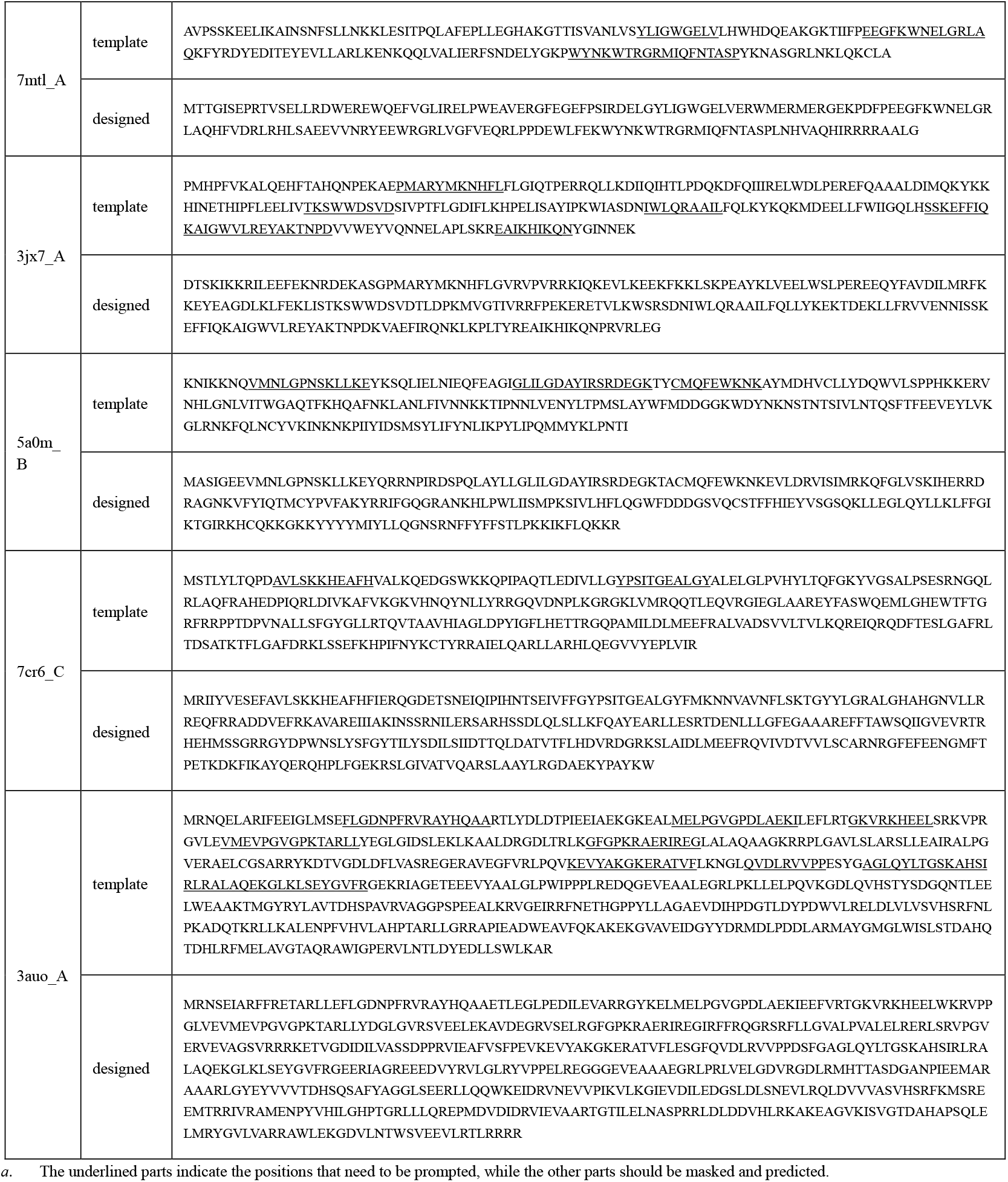
Designed mutants of 10 purified DNA-binding proteins.

**Table S6.**
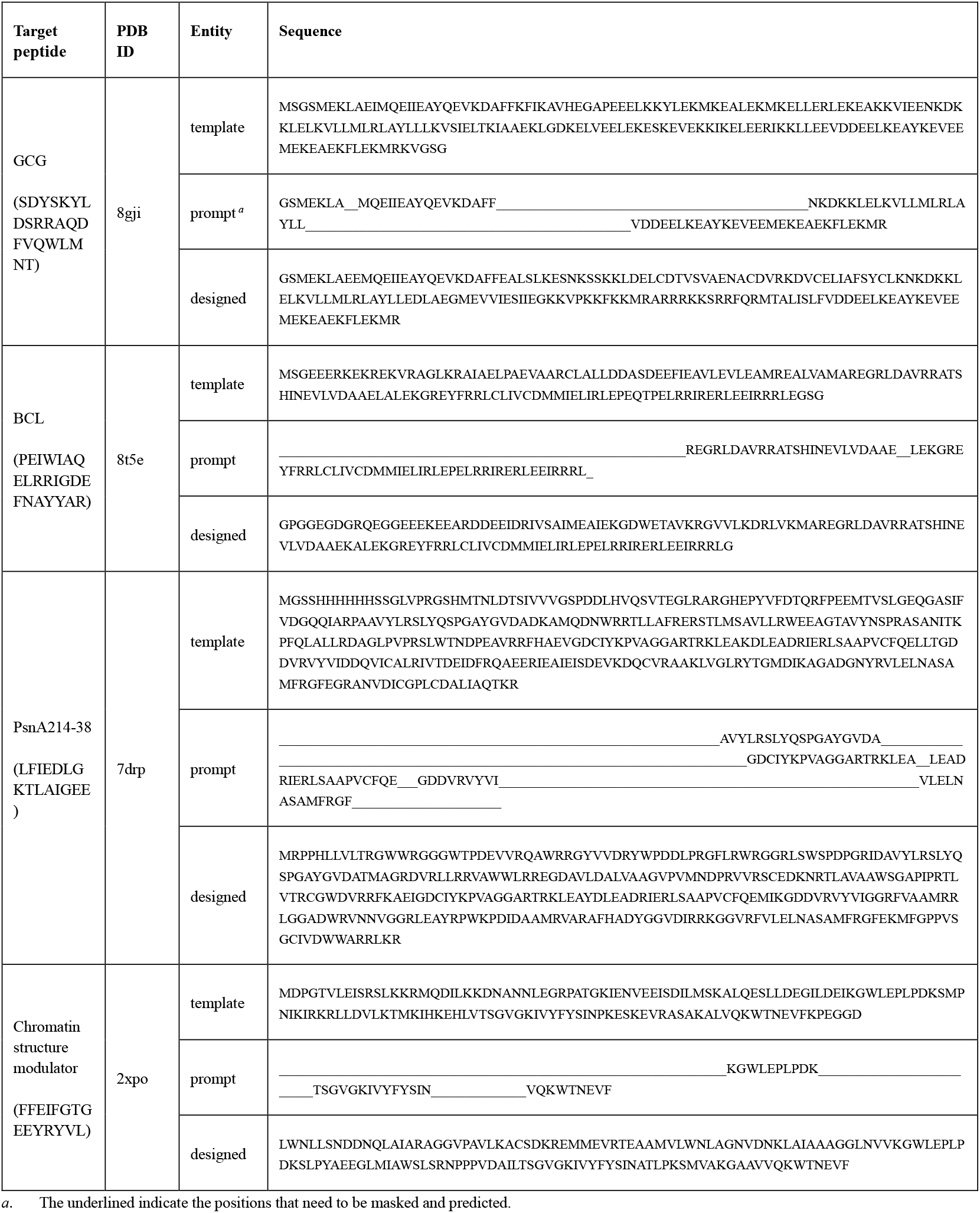
Designed peptide binder sequences.

